# Imaging the pharmacokinetics and therapeutic availability of the bispecific CD3xTRP1 antibody in syngeneic mouse tumor models

**DOI:** 10.1101/2023.06.06.543829

**Authors:** Gerwin G.W. Sandker, Jim Middelburg, Evienne Wilbrink, Janneke Molkenboer-Kuenen, Erik H.J.G. Aarntzen, Thorbald van Hall, Sandra Heskamp

## Abstract

**Background:** CD3 bispecific antibodies (CD3-bsAbs) require binding of both a tumor-associated surface antigen and CD3 for their immunotherapeutic effect. Their efficacy is, therefore, influenced by the absolute tumor uptake and the extracellular dose. To optimize their currently limited efficacy in solid tumors, increased understanding of their pharmacokinetics and *in vivo* internalization is needed.

**Methods:** Here were studied the pharmacokinetics and *in vivo* internalization of CD3xTRP1, a fully murine Fc-inert bsAb, in endogenous TRP1-expressing immunocompetent male C57BL/6J mice bearing TRP1-positive and negative tumors over time. Matching bsAbs lacking TRP1- or CD3-binding capacity served as controls. BsAbs were radiolabeled with ^111^In to investigate their pharmacokinetics, target binding, and biodistribution through SPECT/CT imaging and *ex vivo* biodistribution analyses. Co-injection of ^111^In- and ^125^I-labeled bsAb was performed to investigate the *in vivo* internalization by comparing tissue concentrations of cellular residing ^111^In versus effluxing ^125^I. Anti-tumor therapy effects were evaluated by monitoring tumor growth and immunohistochemistry.

**Results:** SPECT/CT and biodistribution analyses showed that CD3xTRP1 specifically targeted TRP1-positive tumors and CD3-rich lymphoid organ and uptake peaked 24 hours pi (KPC3-TRP1: 37.7±5.3 %ID/g, spleen: 29.0±3.9 %ID/g). Studies with control bsAbs demonstrated that uptake of CD3xTRP1 in TRP1-positive tumors and CD3-rich tissues was primarily receptor-mediated. Together with CD3xTRP1 in the circulation being mainly unattached, this indicates that CD3^+^ T cells are generally not traffickers of CD3-bsAbs to the tumor. Additionally, “antigen-sink” effects by TRP1-expressing melanocytes were not observed. We further demonstrated rapid internalization of CD3xTRP1 in KPC3-TRP1 tumors (24h pi: 54.9±2.3% internalized) and CD3-rich tissues (spleen, 24h pi: 79.7±0.9% internalized). Therapeutic effects by CD3xTRP1 were observed for TRP1-positive tumors and consisted of high tumor influx of CD8^+^ T cells and neutrophils, which corresponded with increased necrosis and growth delay.

**Conclusions:** We show that CD3xTRP1 efficiently targets TRP1-positive tumors and CD3-rich tissues primarily through receptor-mediated targeting. We further demonstrate rapid receptor-mediated internalization of CD3xTRP1 in TRP1-positive tumors and CD3-rich tissues. Even though this significantly decreases the therapeutical available dose, CD3xTRP1 still induced effective anti-tumor T-cell responses and inhibited tumor growth. Together, our data on the pharmacokinetics and mechanism of action of CD3xTRP1 pave the way for further optimization of CD3-bsAb therapies.

**Graphical abstract:** Imaging the pharmacokinetics and therapeutic availability of the bispecific CD3xTRPl antibody in syngeneic mouse tumor models

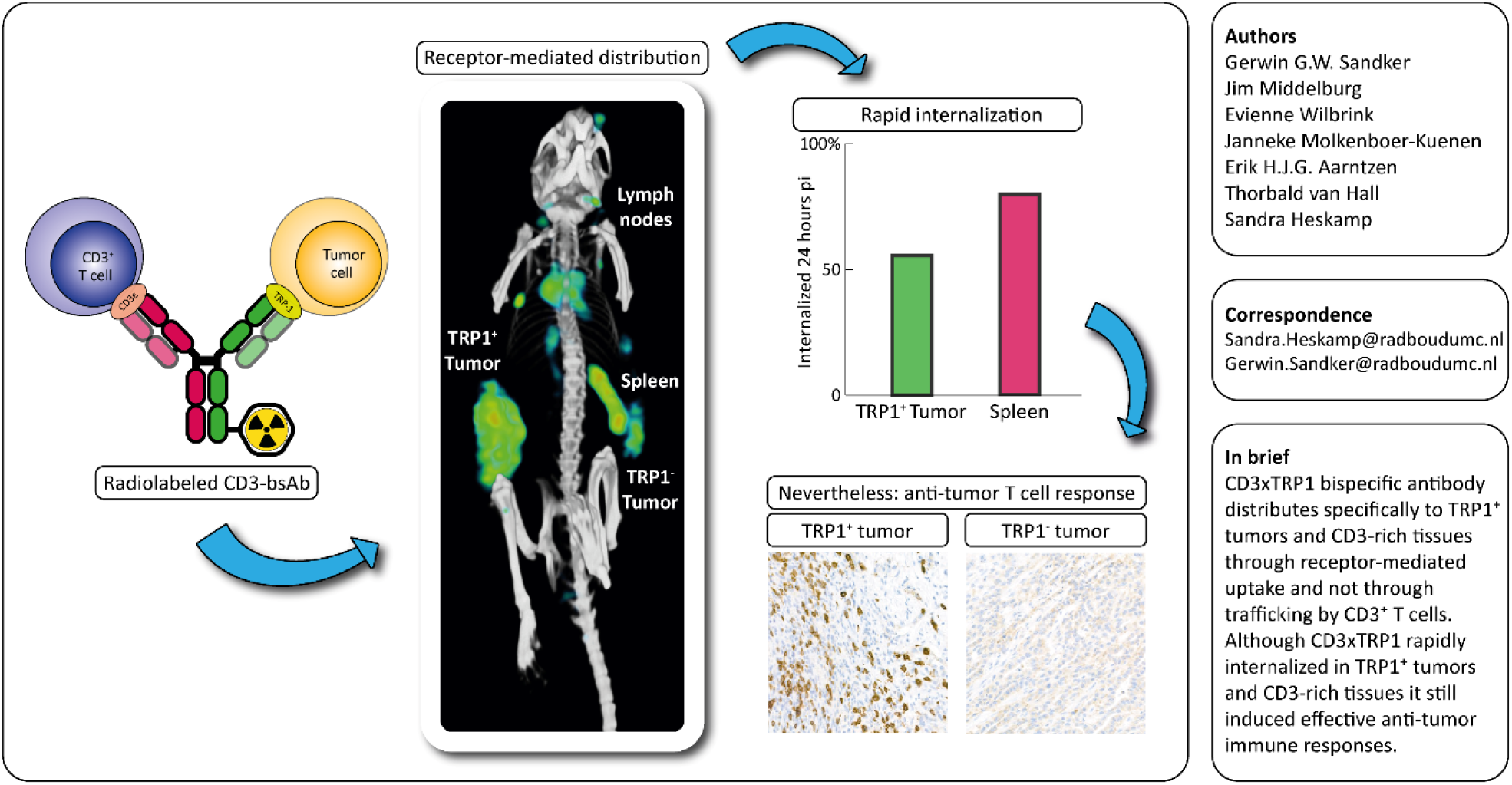

## Introduction

Immunotherapeutic CD3-targeting bispecific antibodies (CD3-bsAbs) show therapeutic potential in a multitude of malignancies [1]. These bsAbs are designed to specifically bind CD3 and a tumor-associated surface antigen (TAA). Upon simultaneous binding to CD3 and the TAA, an immunological synapse is formed, resulting in local T-cell activation and tumor-cell killing independent of TCR specificity, and thus elicits anti-tumor activity even from non-tumor-specific T cells [2].

CD3-bsAbs have shown impressive clinical results for hematological malignancies [3, 4]. In solid cancers, however, their therapeutic efficacy is still limited for several reasons. First, their immunosuppressive tumor microenvironment is characterized by poor infiltration and reduced effector T-cell function [5]. Second, poor vascularization and low or heterogeneous TAA expression may hamper CD3-bsAbs tumor accumulation [6]. Third, expression of TAAs on healthy tissues potentially affects pharmacokinetics (PK) and biodistributions of CD3-bsAbs through the “antigen sink” effect. Furthermore, it increases the potential for adverse effects through “on-target, off-tumor” binding [7]. Thus, factors affecting tumor uptake and PK play an important role CD3-bsAbs’ efficacy and toxicity profiles, especially for solid tumors [8]. Additionally, as the binding of CD3-bsAbs to both the TAA and CD3 on the cell membrane is required to induce the formation of the cytolytic synapse, its extracellular localization is essential for eliciting anti-tumor T-cell responses. Internalization of CD3-bsAbs consequently negates their therapeutic potential. Therefore, thorough understanding of CD3-bsAbs’ PK and *in vivo* cellular internalization is essential to increase efficacy of novel CD3-bsAbs for solid cancers.

Nuclear imaging of radiolabeled CD3-bsAbs allows for evaluating their PK and biodistribution on a full-body scale. Preclinical models enable detailed translational studies to investigate more complex characteristics, such as *in vivo* internalization. As preclinical studies with human CD3-bsAbs are restricted to immunodeficient or artificial models because they lack cross-species specificity, they omit the complex immunological interplay of CD3-bsAbs with their targets. This considerably complicates the translation of results to the clinical setting. Therefore, we investigated the PK, biodistribution, and *in vivo* internalization of a fully murine Fc-inert bsAb targeting murine CD3ε and the melanoma-associated antigen murine tyrosinase-related protein 1 (TRP1, also gp75 or TYRP-1)(CD3xTRP1 bsAb) that has been extensively evaluated for its immunological effects [9, 10], and whose human analogue is being clinically investigated [11].

Here, we report the PK, biodistribution, and *in vivo* internalization of radiolabeled CD3xTRP1 bsAb and control bsAbs in fully syngeneic tumor mouse models with endogenous TRP1 expression. We show that CD3xTRP1 specifically accumulates in TRP1 expressing tumors and CD3-rich tissues. Furthermore, effective anti-tumor immune responses were induced, even though rapid bsAb internalization in TRP1 expressing tumors and secondary lymphoid organs (SLOs) occurred.

## Methods

### Cell culture

KPC3 murine pancreatic ductal adenocarcinoma cells (further KPC3) were transfected to express TRP1 (KPC3-TRP1), whereas B16F10 murine melanoma cells (CRL6475, ATCC) have endogenously TRP1 expression levels [9, 12]. KPC3-TRP1, KPC3, and B16F10 cells were cultured in Iscove′s Modified Dulbecco′s Medium (IMDM, Gibco) supplemented with L-glutamine, 25 mM HEPES, and 8% fetal calf serum (FCS, Sigma-Aldrich Chemie BV). TK-1 murine T-cell lymphoma cells (CRL-2396, ATCC) were cultured in Dulbecco’s Modified Eagle Medium high glucose (DMEM GlutaMAX™, Gibco) supplemented with 1 mM pyruvate (Gibco) and 10% FCS. All cells were cultured at 37°C in a humidified atmosphere with 5% CO2. Cells were tested and found negative for mycoplasma and mouse pathogens.

### bsAbs and radiolabeling

Fc-γ effector-function silenced IgG2a bsAb CD3xTRP1 (2C11 and TA99 arms [13, 14]), and control CD3xMock and TRP1xMock whose TRP1 or CD3-arm was replaced with non-relevant influenzavirus-specific “Mock” arms (B12 [15]), were previously produced and provided by Genmab [10]. BsAbs were conjugated with a 15-fold molar excess of isothiocyanatobenzyl-diethylenetriaminepentaacetic acid (ICT-DTPA, p-SCN-Bn-DTPA, Macrocyclics) in 0.1 M NaHCO3 (pH 9.5), for 1 hour at RT. Subsequently, unreacted ITC-DTPA was removed by dialysis against 0.25 M ammonium acetate (NH4Ac) pH 5.4–5.5 (metal free) supplemented with 2 g/L Chelex (Bio-Rad) using 20,000 molecular weight cut-off dialysis cassettes (Slide-a-Lyzer, Thermofisher). Radiochemical analysis showed conjugation ratios of 1.62, 1.59, and 1.37 ITC-DTPA/antibody for CD3xTRP1, CD3xMock, and TRP1xMock, respectively.

DTPA-bsAbs were radiolabeled with ^111^InCl_3_ (Curium) in 0.5 M MES buffer (pH 5.4) for 15-20 minutes at RT. Next, unincorporated ^111^In was chelated by adding ethylenediaminetetraacetic (EDTA) in PBS to a final 5 mM concentration. Iodine-125 (^125^I) labeling was performed by incubating DTPA-bsAbs in iodogen coated Eppendorf tubes with Na^125^I (10^−5^ M NaOH, Perkin Elmer) in 0.5 M phosphate buffer (pH 7.4) with the total reaction volume adjusted to 100 µL with H2O, for 10 minutes at RT. The reaction was quenched with a saturated tyrosine solution. Labeling efficiencies were evaluated with instant thin-layer chromatography (iTLC), using silica gel chromatography strips (Agilent Technologies) and 0.1 M, pH 6.0 citrate (Sigma-Aldrich) as running buffer. The iTLC activity profiles were analyzed using photosensitive plates (Fuji MS, Cytiva), a phosphor imager (Typhoon FLA 7000, GE), and AIDA software v4.21.033 (Raytest, Straubenhardt). Labeling efficiencies were >95% for *in vivo* and >90% for *in vitro* experiments. For *in vivo* experiments, the reaction mixture was adjusted to pH 7.4 with 10x PBS (pH 7.4). Unlabeled bsAb and sterile pyrogen-free PBS was added to reach a final concentration of 62.5 µg/mL.

### *In vitro* characterization

We evaluated whether DTPA-conjugation and radiolabeling affected CD3- or TRP1-binding of CD3xTRP1 and control bsAbs with flow cytometry, Lactate Dehydrogenase (LDH) cytotoxicity, and *in vitro* binding assays.

### Flow cytometry

KPC3 and TRP1-positive B16F10 cells were stained with 10 µg/mL unconjugated CD3xTRP1 or DTPA-conjugated CD3xTRP1 in PBS supplemented with 0.5% bovine serum albumin (BSA, Sigma) and 0.002% sodium azide (FACS buffer) for 20 minutes at 4°C. Next, cells were washed with FACS buffer and stained with goat-anti-mouse IgG-Alexa Fluor647 (1:400, Biolegend) in FACS buffer for 20 minutes at 4°C. Finally, cells were resuspended in FACS buffer, measured on a flow cytometer (LSRFortessa™ X-20, BD bioscience), and analyzed with FlowJo v10.8.1 (Treestar).

### LDH cytotoxicity assay

Proliferation of KPC3 and KPC3-TRP1 was arrested by irradiation with 6000 Rad (Gammacell 1000 Elite, Best Theratronics). Subsequently, 30,000 tumor cells/well were seeded in 96-wells plates and allowed to adhere for 4 hours. T cells were isolated from spleens of naïve C57BL/6J mice using nylon wool fiber columns to remove B cells. Then, 300,000 T cells/well and either non-conjugated or DTPA-conjugated CD3xTRP1 bsAbs was added and incubated for two days at 37ºC. Tumor cell killing was assessed using the CyQUANT™ LDH cytotoxicity assay (Thermo Fisher) following the manufacturers protocol. Absorbance was measured on a Spectramax iD5 plate-reader (Molecular Devices). The percentage of cytotoxicity was calculated, after correcting for absorbance of medium only and tumor cell only controls, from the absorbance of the treated samples and Triton100-lysed maximum kill control samples.

### *In vitro* binding assay

KPC3-TRP1 (1.4×10^6^ cells/mL), KPC3 (1.4×10^6^ cells/mL), and B16F10 (1.4×10^6^ cells/mL) cells were resuspended in IMDM/0.5%BSA, and TK-1 (4.0×10^6^ cells/mL) cells in DMEM/0.5%BSA, and incubated with equimolar concentrations (0.75 pM) of [^111^In]In-CD3xTRP1 (0.85 kBq) or [^125^I]I-CD3xTRP1 (0.7 kBq) for 3.5 hours at 37ºC. Cells were washed twice and cell-associated activity was subsequently measured in a well-type gamma-counter (Wallac 2480 wizard, Perkin Elmer). In a separate binding assay equimolar concentrations (1.97 pM) of [^111^In]In-CD3xMock (0.76 kBq) and [^111^In]In-TRP1xMock (0.75 kBq) were incubated with KPC3-TRP1 (0.8×10^6^ cells/mL), KPC3 (2.6×10^6^ cells/mL), or TK-1 (8.5×10^6^ cells/mL) in RPMI-1640/0.5%BSA (Gibco) for 40 minutes at 37ºC. Cell-associated activity was measured as described above.

### Animal studies

Male C57BL/6J mice (n=78, 6–8 weeks, Charles River, Germany) were housed under specific-pathogen-free conditions in individually ventilated cages with a filter top (Green line IVC, Tecniplast), in the presence of cage enrichment with food and water ad libitum. For tumor inoculation, mice were injected subcutaneously with 200 µL PBS/0.1%BSA containing; 1) 0.8×10^5^ KPC3-TRP1 cells (right flank) and 0.8×10^5^ KPC3 cells (left flank), or 2) 0.8×10^5^ B16F10 cells (right flank). Tumor size was measured using a caliper twice weekly. KPC3 and KPC3-TRP1 tumors-bearing mice were block-randomized based on tumor size and injected with radiolabeled bsAbs 17 days after inoculation. B16F10 tumor-bearing mice received [^111^In]In-CD3xTRP1 eight days post inoculation. Confounding effects of cage allocation were prevented by pooling mice from different groups. Humane endpoints were based on the general level of discomfort and tumor sizes; animal healthy was checked daily. All *in vivo* experiments and corresponding protocols were approved by the Animal Welfare Body of the Radboud University, Nijmegen, and the Central Authority for Scientific Procedures on Animals (AVD1030020209645) and were performed in accordance with the principles stated by the Dutch Act on Animal Experiments (2014). Researchers and biotechnicians were blinded for group allocation.

### *Ex vivo* biodistribution analyses

To assess the pharmacokinetics, biodistribution, and therapeutic availability of CD3xTRP1 we performed *ex vivo* biodistribution experiments with ^111^In- and ^125^I-labeled CD3xTRP1, CD3xMock, and TRP1xMock in KPC3 and KPC3-TRP1 tumors-bearing mice at several time points. Mice (n=7/group) were injected intraperitoneally in the right caudal abdomen with 200 µL PBS/0.1%BSA containing 12.5 µg of a 1:1 mixture of ^111^In- and ^125^I-labeled; a) CD3xTRP1 ([^111^In]In and [^125^I]I: 0.43 and 0.17 MBq, (n=21)), b) CD3xMock (0.44 and 0.17 MBq, (n=21)), or c) TRP1xMock (0.43 and 0.17 MBq, (n=21)). At 24, 72, or 168 hours post injection (pi), mice were euthanized by CO2/O2 asphyxiation, blood was collected via heart puncture and relevant tissues (blood, muscle, shaven pigmented skin, shaven non-pigmented skin, right and left inguinal lymph nodes, tumors, spleen, thymus, bone marrow, bone, pancreas, duodenum, heart, lung, kidney, liver, stomach, colon, brown adipose tissue, prostate) were collected for biodistribution analysis. To separate the serum and cell pellet, blood was stored in an Eppendorf tube for 30 minutes at RT, followed by centrifugation at 2,000 g for 10 minutes at 4ºC. Tissue and blood samples were weighed (XPE105DR, Mettler Toledo) and radioactivity concentrations were measured with a well-type gamma-counter. Aliquots of injection fluid served as reference controls. Tissue accumulation of [^111^In]In-bsAb and [^125^I]I-bsAb was calculated as a percentage of the injected dose and normalized for tissue weight (percentage injected dose per gram (%ID/g)). To evaluate the absolute accumulated dose in the tumors, spleen, and lymph nodes, we calculated the percentage of the injected dose (%ID) accumulated in the entire tissue. *In vivo* internalization of bsAbs was investigated by evaluating ^125^I-to-^111^In ratios of tissue uptake concentrations; upon internalization and subsequent degradation ^125^I effluxes from the cell [16], whereas ^111^In resides intracellularly [17]. Tumor tissues were fixated in PBS/4%formalin for immunohistochemical (IHC) analysis.

### SPECT/CT imaging

To longitudinally visualize the *in vivo* biodistribution of CD3xTRP1 and control bsAbs in KPC3 and KPC3-TRP1 tumors-bearing and B16F10 tumor-bearing mice, we performed SPECT/CT imaging at various time points. KPC3 and KPC3-TRP1 tumor-bearing mice (n=3/group) were injected intraperitoneally in the right caudal abdomen with 12.5 µg [^111^In]In-CD3xTRP1, [^111^In]In-CD3xMock, or [^111^In]In-TRP1xMock (15.4–17.7 MBq). SPECT/CT (U-SPECT II/CT, MILabs) images were acquired at 6, 24, and 72 hours pi (acquisition times: 15, 25, and 35 minutes, respectively), under inhalation anesthesia (2%isoflurane in air) on a heated bed (38°C), and 168 hours pi following CO2/O2 asphyxiation (acquisition time: 55 min), using a 1.0 mm diameter pinhole mouse collimator, and CT parameters of 160 µm spatial resolution, 615 µA, and 65 kV. Data were reconstructed with MILabs software v2.04 using the following settings: energy windows at 171 keV (154–188 keV) and 245 keV (220–270 keV), 1 iteration, 16 subsets and a 0.2 mm voxel size. We used Inveon Research Workplace software v4.2) to create maximum-intensity projections (MIP) after applying a gaussian 2x2x2 voxel filter. In a separate experiment, B16F10 tumor-bearing mice (n=6) were injected intraperitoneally in the right caudal abdomen with 12.5 µg [^111^In]In-CD3xTRP1 (14.2–14.5 MBq), followed by SPECT/CT imaging at 24 hours pi under inhalation anesthesia (2% isoflurane in air) and 72 hours pi following CO2/O2 asphyxiation, using the same acquisition parameters as described above. After SPECT/CT, *ex vivo* biodistribution analyses (without serum analysis) were performed as described above.

### Immunohistochemistry

Consecutive KPC3 and KPC3-TRP1 tumor sections (5 µm) were evaluated for CD3, CD8, Ly6G, F4/80, PD-L1, and H&E. The staining-procedure for H&E, Ly6G, F4/80, PD-L1 were described previously [18]. For CD8 and CD3, sections were deparaffinized and rehydrated, followed by antigen retrieval in 1x Tris, Borate, EDTA buffer (Amresco/VWR, pH 8.3) containing 0.05% Tween-20 at 96°C for 10 minutes. Then, endogenous peroxidases, endogenous biotin, and non-specific Ig binding were blocked using 3% H2O2, a biotin/avidin blocking set (SP-2001, Vector), and normal goat serum (5095, Bodinco), respectively. Primary antibody rabbit-anti-mouse CD8alpha (1:4,000, ab209775, Abcam) was incubated overnight at 4°C, and rabbit-anti-mouse CD3 (1:400, ab16669, Abcam) for 60 minutes at RT. Next, secondary antibody goat-anti-rabbit/biotin antibody (1:200, BA-1000, Vector) was incubated for 30 minutes at RT. Thereafter, for signal amplification, sections were incubated with horseradish peroxidase-Biotin/Streptavidin Complex (1:100, PK-6100, Vectastain) for 30 minutes at RT. Finally, sections were incubated with bright DAB (B-500, Immunologic) for 8 minutes, counterstained with hematoxylin (Klinipath/VWR), dehydrated, and mounted with a cover slip using permount (Fisher). Sections were analyzed with blinding for group allocation. Representative tumors sections were scanned at 20x magnification using a Pannoramic 1000 (0.242535x0.242647 µm/pixel, 3DHISCTECH). Images were created from representative tumor areas using Caseviewer v2.3. Intratumoral CD3^+^ and CD8^+^ T-cell presence was quantified on images (n=6/section) at 10x magnification in Fiji v1.53t using; color deconvolution (H DAB), auto threshold “Yen”, and analyze particles (size=10 to 400) to calculate the %DAB-stained area.

### Statistical analyses

The required number of animals was determined by *a priori* sample size calculations based on previously obtained tumor uptake values. 11 mice were excluded from analysis as; 1) one B16F10 tumor did not develop, 2) one B16F10 tumor-bearing and nine KPC3 and KPC3-TRP1 tumors-bearing mice from various groups were misinjected intraperitoneally, which was posteriorly defined as an [^111^In]In-bsAb blood level ≤1/5^th^ of the groups mean at time of dissection. All results are presented as mean ± standard deviation (sd), unless stated otherwise. Differences in [^111^In]In-bsAbs’ blood concentrations and tissue uptake was tested for significance using two-way ANOVAs with Bonferroni *post hoc* tests. Differences in KPC3 and KPC3-TRP1^+^ tumor volumes, and increases of intratumoral CD3^+^ and CD8^+^ T cells of CD3xTRP1 or control treated mice were tested for statistical significance using one-way ANOVAs with Bonferroni *post hoc* tests and two-tailed t-tests, respectively. Statistical significance was defined as p-value ≤0.05 and were performed with GraphPad Prism v9.5.

## Results

### CD3xTRP1 target binding is unaffected by radiochemical procedures

To assess whether bsAb conjugation with the (radio)metal chelator DTPA and/or radiolabeling affected CD3- or TRP1-binding, we performed flow cytometry, LDH cytotoxicity, and *in vitro* binding assays. Flow-cytometric analysis of equimolar concentrations of unconjugated CD3xTRP1 and DTPA-CD3xTRP1 showed similar binding to TRP1-expressing B16F10 cells, while no binding was observed to parental KPC3 cells (Figure 1A/B). In addition, the LDH cell killing assay showed equal induction of cytotoxicity of naïve splenic T cells to KPC3-TRP1 cells by DTPA-conjugated CD3xTRP1 and unconjugated CD3xTRP1 at all tested doses (Figure 1C). Moreover, both ^111^In- and ^125^I-labeled DTPA-CD3xTRP1 showed specific binding to TRP1 and CD3, which was not significantly different from each other (Figure 1D). Target binding of [^111^In]In-CD3xMock to CD3 and [^111^In]In-TRP1xMock to TRP1 was also confirmed (Figure 1E). Taken together, ITC-DTPA conjugation and radiolabeling of CD3xTRP1 did not interfere with target binding and T-cell activation.

**Figure 1.**
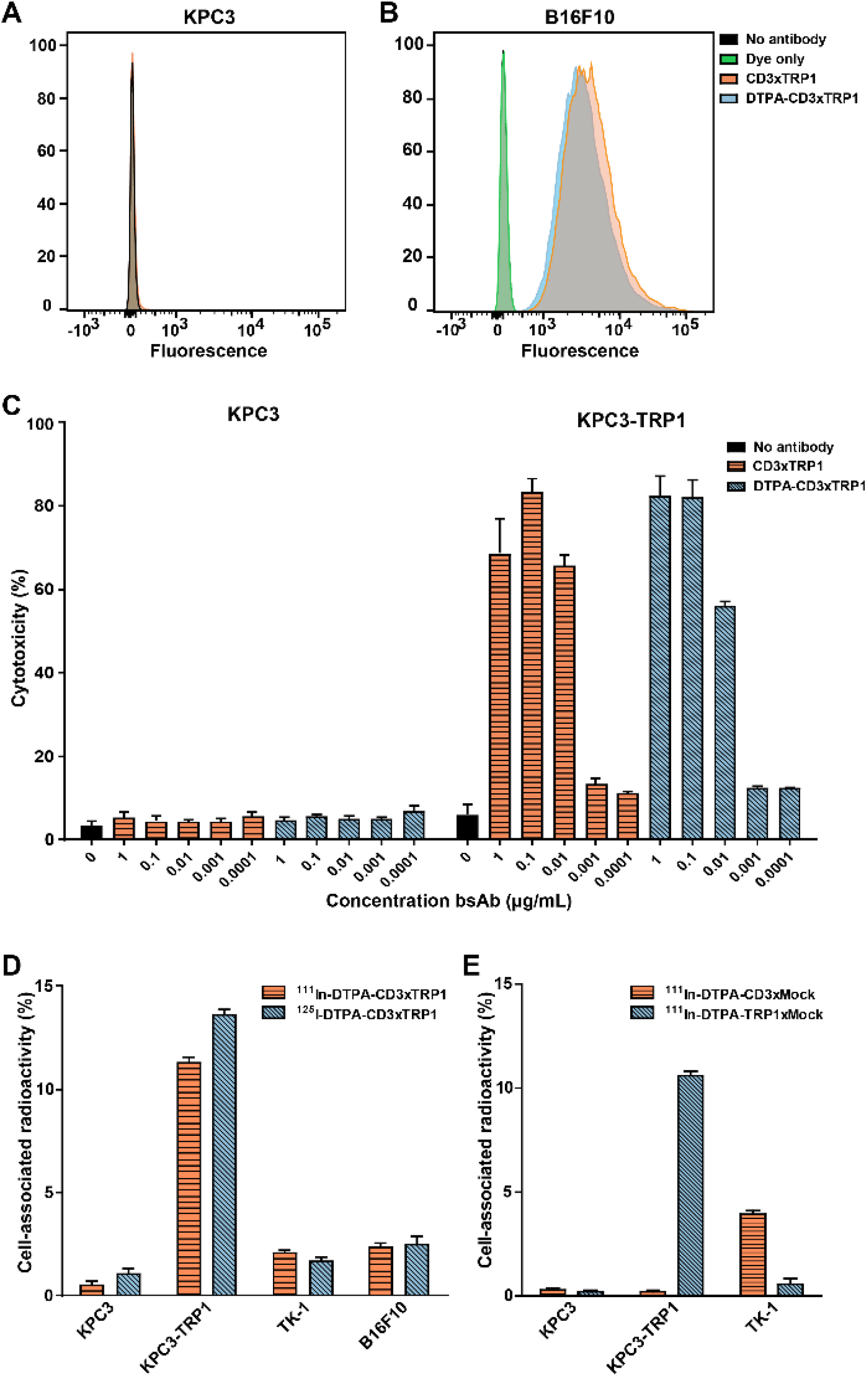
*In vitro* evaluation of cell binding and cytotoxicity of CD3xTRP1 after DTPA-conjugation and radiolabeling. **(A/B)** Flow cytometry analysis of unconjugated and DTPA-conjugated CD3xTRP1 target binding on KPC3 and TRP1 positive B16F10 cells. **(C)** LDH cytotoxicity assay on KPC3 and KPC3-TRP1 cells with unconjugated or DTPA-conjugated CD3xTRP1. **(D)** Binding assays of [^111^In]In-CD3xTRP1 and [^125^I]I-CD3xTRP1 on KPC3, KPC3-TRP1, CD3-positive TK-1, and TRP1-positive B16F10 cells. **(E)** Binding assays of [^111^In]In-CD3xMock and [^111^In]In-TRP1xMock on KPC3, KPC3-TRP1, and CD3^pos^ TK-1 cell lines.

### CD3xTRP1 distribution is mediated via both the CD3- and TRP1-binding arm

We evaluated *in vivo* target specificity of the CD3- and TRP1-binding arms with SPECT/CT and *ex vivo* biodistribution studies using ^111^In-labeled CD3xTRP1 and control bsAbs lacking either the CD3- or TRP1-binding arm. SPECT/CT revealed [^111^In]In-CD3xTRP1 accumulation in CD3-rich tissues and KPC3-TRP1 tumors and not in parental KPC3 tumors. [^111^In]In-CD3xMock merely distributed to CD3-rich tissues and not to KPC3-TRP1 tumors (Figures 2 and S1), while [^111^In]In-TRP1xMock uptake was only observed in KPC3-TRP1 tumors. These results indicate that CD3xTRP1 tissue uptake is target mediated. Additionally, SPECT/CT imaging showed accumulation of all three bsAbs in the peri- and extratumoral region extending caudally towards the hip of both KPC3-TRP1 and KPC3 tumors (Figures 2 and 3), indicating a non-CD3 and non-TRP1 mediated process. Moreover, in B16F10 tumor-bearing mice this accumulation was not observed, suggesting a tumor model specific process (Figure 3A/B).

**Figure 2.**
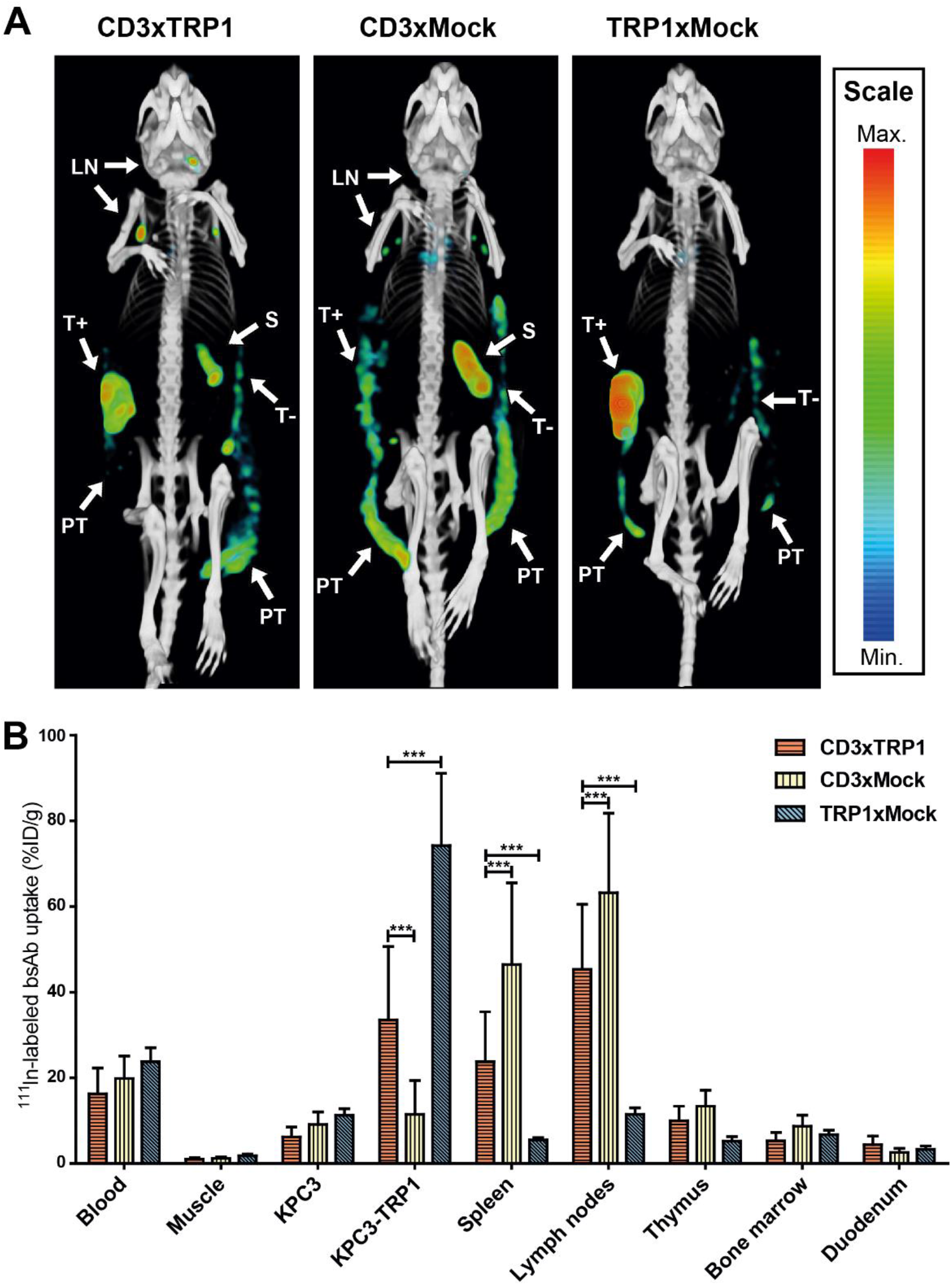
*In vivo* biodistribution of CD3xTRP1, CD3xMock, and TRP1xMock. MicroSPECT/CT and *ex vivo* biodistribution analysis of 12.5 µg [^111^In]In-CD3xTRP1, [^111^In]In-CD3xMock, and [^111^In]In-TRP1xMock in C57BL/6J mice bearing KPC3 and KPC3-TRP1 tumors on contralateral flanks, 72 hours post intraperitoneal injection. (**A)** *In vivo* biodistribution of [^111^In]In-bsAbs is visualized with MIPs generated from microSPECT/CT data using the same thresholds at time point. Indicated tissues with [^111^In]In-bsAbs uptake are: KPC3-TRP1 tumor (T+), KPC3 tumor (T-), spleen (S), lymph nodes (LN), and peri- and extratumoral accumulation (PT). **(B)** Biodistribution of [^111^In]In-CD3xTRP1, [^111^In]In-CD3xMock, and [^111^In]In-TRP1xMock in KPC3 and KPC3-TRP1^+^ tumors-bearing mice at 72 hours pi. Only relevant tissues are depicted. Tissue uptake is presented as %ID/g, differences in uptake were tested for significance using two-way ANOVAs with a Bonferroni *post hoc* test (***=p<0.001).

**Figure 3.**
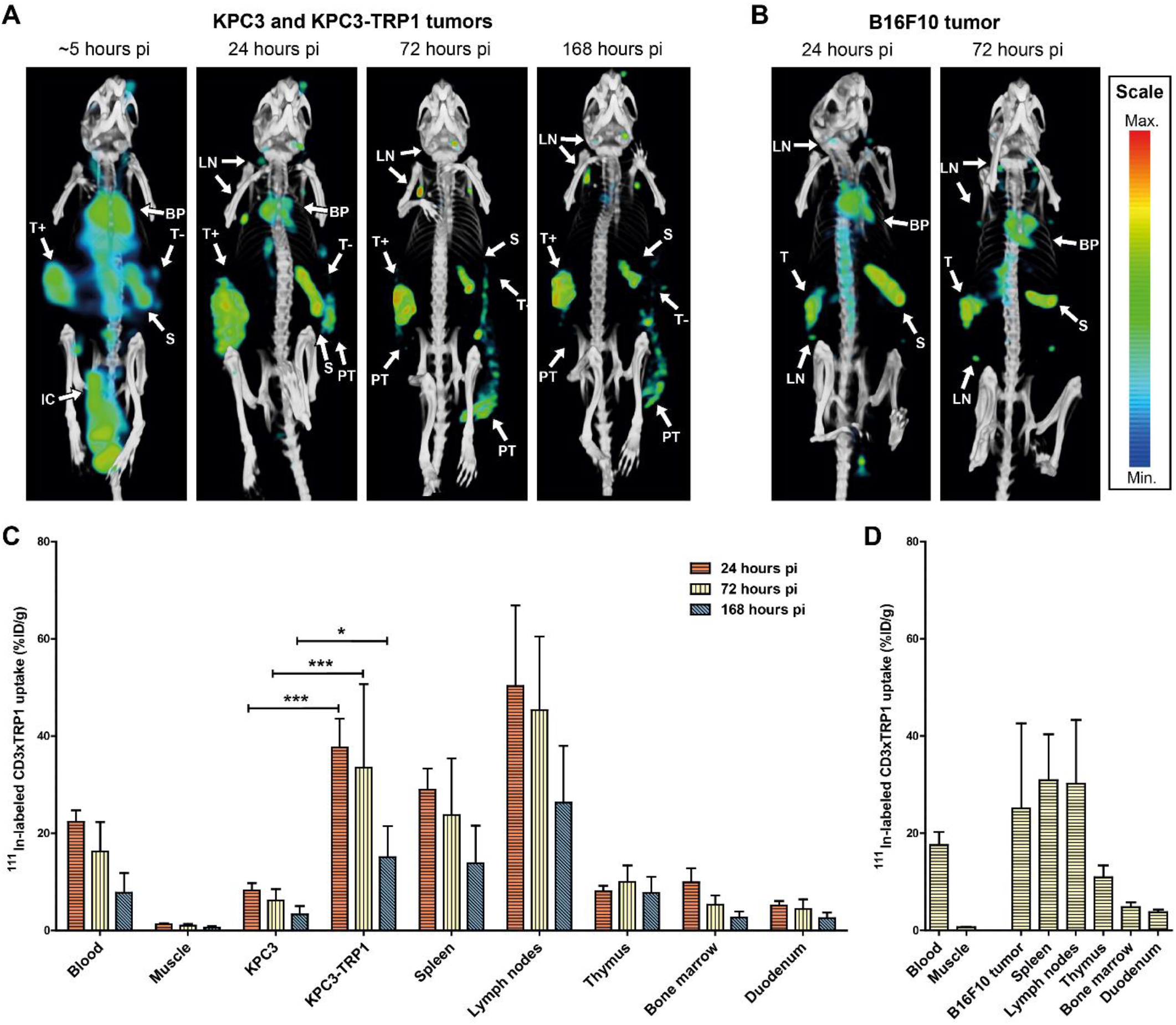
*In vivo* biodistribution of CD3xTRP1 over time. MicroSPECT/CT and *ex vivo* biodistribution analyses of 12.5 µg [^111^In]In-CD3xTRP1 in C57BL/6J mice bearing either; 1) KPC3-TRP1 and KPC3 tumors on contralateral flanks at ∼5, 24, 72, and 168 hours post intraperitoneal injection, or 2) B16F10 tumors at 24 and 72 hours post intraperitoneal injection. **(A)** Representative MIPs generated from microSPECT/CT data using the same thresholds for the distinct time points visualizing tissue uptake of [^111^In]In-CD3xTRP1 in KPC3 tumor bearing mice. **(B)** Representative MIPs generated from microSPECT/CT data using the same thresholds at time point visualizing tissue uptake of [^111^In]In-CD3xTRP1 in B16F10 tumor bearing mice. **(C)** Biodistribution of [^111^In]In-CD3xTRP1 in KPC3 and KPC3-TRP1^+^ tumors-bearing mice at 24, 72, and 168 hours pi. **(D)** Biodistribution of [^111^In]In-CD3xTRP1 in B16F10 tumor-bearing mice at 72 hours pi. Indicated tissues with [^111^In]In-CD3xTRP1 uptake are: blood pool (BP), intraperitoneal cavity (IC), KPC3-TRP1 tumor (T+), KPC3 tumor (T-), B16F10 tumor (T), spleen (S), lymph nodes (LN), and peri- and extratumoral accumulation (PT). Tissue uptake in panel C and D is presented as %ID/g, differences in uptake were tested for significance using two-way ANOVAs with a Bonferroni *post hoc* test (*=p<0.05 and ***=p<0.001).

Biodistribution analyses quantitatively confirmed these results; uptake of [^111^In]In-TRP1xMock in KPC3-TRP^pos^ tumors was significantly higher compared with [^111^In]In-CD3xTRP1 (72 hours pi: 74.2±15.4 vs 33.5±15.4 %ID/g, p<0.001), whereas its uptake in CD3-rich tissues, such as the spleen, was significantly lower (Figure 2, table S1). Inversely, uptake of [^111^In]In-CD3xMock in KPC3-TRP1 tumors was significantly lower compared with bsAbs with a TRP binding arm, while uptake in CD3-rich organs was significantly higher (spleen 72 hours pi, CD3xMock vs CD3xTRP1: 46.4±17.7 vs 23.8±10.4 %ID/g, p<0.001). Additionally, this suggests that KPC3-TRP1 and SLO uptake of CD3xTRP1 was not saturated. Blood levels and marker negative tissue uptake were unaffected by removal of CD3- or TRP1-binding capacity. In pigmented and non-pigmented skin [^111^In]In-CD3xTRP1 uptake was not statistically significant different. Additionally, it was comparable to skin uptake of CD3xMock and TRP1xMock (table S1). This indicated no pronounced specific uptake of CD3xTRP1 in healthy pigmented skin. In summary, SPECT/CT imaging and biodistribution analyses show CD3- and TRP1-specificity of CD3xTRP1.

### CD3xTRP1 efficiently targets tumors expressing TRP1

To investigate the *in vivo* distribution of CD3xTRP1 over time, we evaluated the pharmacokinetics of [^111^In]In-CD3xTRP1 in C57BL/6J mice bearing: 1) KPC3 and KPC3-TRP1 tumors, and 2) B16F10 tumors. CD3xTRP1 blood levels were highest at 24 hours pi and decreased gradually over time (Figure 3). KPC3-TRP1 tumor uptake was highest at 24 hours pi (37.7±5.3 %ID/g) and steadily decreased thereafter (168 hours: 15.1±5.8 %ID/g, p<0.001), but was significantly higher compared with KPC3 tumor uptake (168 hours: 3.3±1.5 %ID/g, p<0.05). Spleen, lymph node, and bone marrow showed similar trends with CD3xTRP1 uptake peaking 24 hours pi, whereas thymic uptake peaked at 72 hours pi. For a comprehensive overview of the biodistributions see supplemental table S1. As tumor growth continued after bsAb injection, which decreases the weight-corrected uptake concentrations, we also evaluated the absolute tumor uptake in %ID. We observed that the absolute KPC3-TRP1 tumor uptake of CD3xTRP1 was constant over time. Thus, the decreased tumor uptake concentrations were primarily a result of dilution through tumor growth accompanied with low additional CD3xTRP1 accumulation (table S2). SPECT/CT imaging results corresponded with the *ex vivo* biodistribution analyses (Figure 3A/C). Additionally, SPECT/CT imaging showed rapid [^111^In]In-bsAb absorption into the circulation, and uptake in the spleen and KPC3-TRP1 tumor within the first 4 hours following intraperitoneal injection. Lymph node uptake became discernible at 24 hours pi.

These findings were confirmed in a second tumor model B16F10, whose endogenous TRP1 expression level is lower compared with KPC3-TRP^pos^, which expresses TRP1 through transfection [9]. SPECT/CT and *ex vivo* biodistribution analysis revealed similar pharmacokinetics of [^111^In]In-CD3xTRP1 (Figure 3B/D), but B16F10 tumor uptake was lower than KPC3-TRP1 tumor uptake, although not statistically significant (25.1±15.1 vs 33.5±15.4 %ID/g, p=ns) (table S1). Inguinal lymph node uptake was significantly lower in B16F10 tumor-bearing mice compared with KPC3 tumors-bearing mice (30.2±11.3 vs 45.3±13.6 %ID/g, p<0.01).

To evaluate whether CD3xTRP1 in circulation was bound to circulating CD3^+^ cells, we determined the localization of [^111^In]In-CD3xTRP1 in serum and clotted fractions of blood samples over time. All serum fractions contained ∼80% of the radiolabel, suggesting that CD3xTRP1 was generally unbound and continuously available for target binding (table S5).

### CD3xTRP1 undergoes target-mediated internalization

To investigate the availability of tumor- and tissue-accumulated CD3xTRP1 for interaction with CD3, we assessed the internalization of CD3xTRP1 by co-administering ^125^I- and ^111^In-labeled bsAbs [19]. Upon bsAb internalization, ^125^I is released from the cell, while ^111^In is retained intracellularly. Therefore, the ^125^I-to-^111^In ratio provides an estimation of how much accumulated bsAb was internalized. In KPC3-TRP1 tumors, [^125^I]I-CD3xTRP1 uptake (17.0±2.7 %ID/g) was significantly lower than [^111^In]In-CD3xTRP1 uptake at 24 hour pi (37.7±5.3 %ID/g, p<0.001), resulting in an ^125^I/^111^In ratio of 0.45±0.02, meaning that at least 55% of the accumulated CD3xTRP1 was internalized (Figure 4, table S3/4). Significant internalization continued thereafter resulting in at least 80% of KPC3-TRP1 tumor-accumulated CD3xTRP1 to be unavailable for CD3 interaction at both 72 and 168 hours pi (^125^I/^111^In-ratios: 0.22 ± 0.03 and 0.21±0.03). In CD3-rich tissues internalization occurred even faster with 80% of spleen and lymph node accumulated CD3xTRP1 being internalized at 24 hours pi (spleen ^125^I/^111^In-ratio: 0.20±0.01). In the blood and marker negative tissues the ^125^I/^111^In ratio for CD3xTRP1 remained consistent over time, indicating no internalization.

**Figure 4.**
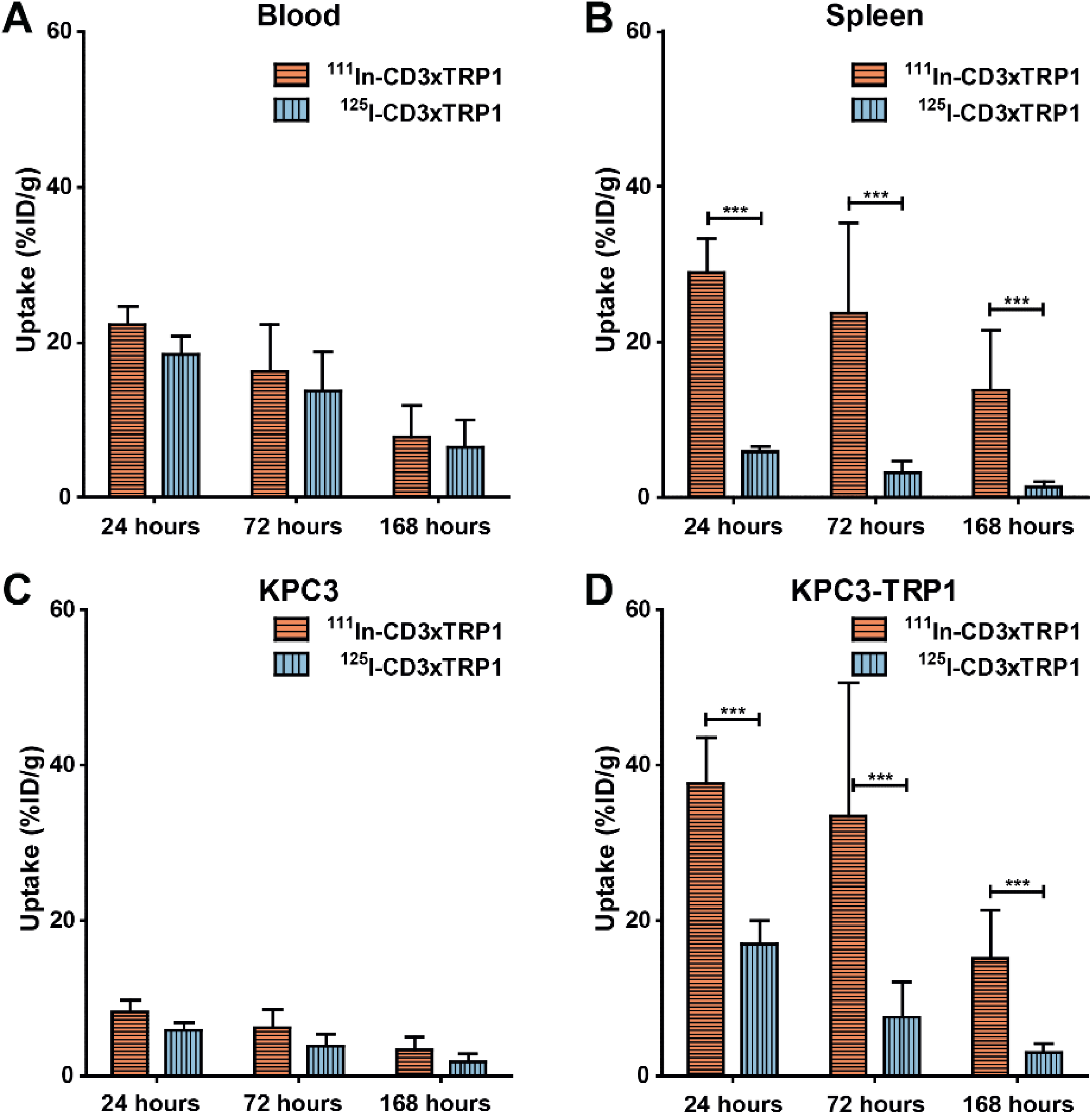
Uptake of [^111^In]In-CD3xTRP1 and [^125^I]I -CD3xTRP1 in relevant tissues over time. Tissue accumulation of intracellular residing [^111^In]In-CD3xTRP1 and cellular effluxing [^125^I]I-CD3xTRP1 in C57BL/6J mice bearing KPC3-TRP1 and KPC3 tumors that received 12.5 µg of a 1:1 mixture of ^111^In- and ^125^I-labeled CD3xTRP1, at 24, 72, and 168 hours post intraperitoneal injection. Blood **(A)**, spleen **(B)**, KPC3 **(C)**, and KPC3-TRP1 **(D)** tumor uptake are given as %ID/g, differences in uptake were tested for significance using two-way ANOVAs with a Bonferroni *post hoc* test (***=p<0.001).

In accordance with its targets, CD3xMock internalization in CD3-rich tissues was similarly high as CD3xTRP1 (table S4), whereas the ^125^I/^111^In ratios of CD3xMock were comparable in KPC3 and KPC3-TRP1 tumors. Likewise, TRP1xMock internalization in KPC3-TRP1 tumors was similarly high when compared to CD3xTRP1, but considerably lower in CD3-rich tissues. This indicated that internalization of CD3xTRP1 in KPC3-TRP1 tumors and CD3-rich tissues were mostly TRP1- or CD3-mediated. Non-target mediated internalization, however, also occurred as the ^125^I/^111^In ratios of CD3xTRP1 and control bsAbs decreased over time in TRP1-negative KPC3 tumors. Additionally, the ^125^I/^111^In ratio of TRP1xMock decreased over time in SLOs, indicating internalization through non-specific processes (table S4). In summary, CD3xTRP1 rapidly internalized *in vivo* in KPC3-TRP1 and CD3-rich tissues, the observed internalization was mostly target-mediated.

### CD3xTRP1 reduces KPC3-TRP1 tumor growth and induces immune-cell infiltration

We assessed the therapy effects of a single injection of radiolabeled CD3xTRP1 and control bsAbs by measuring tumor sizes and evaluating the tumor microenvironment for CD3, CD8, Ly6G, F4/80, and PD-L1. KPC3-TRP1 tumors were significantly smaller in mice treated with CD3xTRP1 compared with TRP1xMock at 168 hours pi (mean±sd: 408±170 vs 751±246 mm^3^, p<0.05) (figure 5A), whereas KPC3 tumor volumes showed no statistically significant differences. Immunohistochemistry revealed substantial increases in necrotic tumor surface area in CD3xTRP1 treated KPC3-TRP1 tumors. This corresponded with increased CD3^+^ and CD8^+^ T-cell numbers in KPC3-TRP1 tumors, as determined by the proportion of CD3- or CD8-stained tumor area, which was most pronounced at 72 hours pi (Figure 5B/C, KPC3 vs KPC3-TRP1: 0.11±0.02 vs 5.45±0.95 %CD3-DAB-stained area, p<0.01). Furthermore, we observed an increase of neutrophils (Ly6G^pos^) and a shift from a heterogeneous to more homogeneous intratumoral neutrophil distribution in vital KPC3-TRP1 tumor regions (figure S2). Additionally, high numbers of Ly6G^pos^ neutrophils and PD-L1^pos^ cells were observed in necrotic areas. Moreover, F4/80^high^ macrophages surrounded the tumor tissue, while tumor infiltrated macrophages appeared to be F4/80^low^. The number and intratumoral distribution of both F4/80^high^ and F4/80^low^ macrophages did not change over time. Finally, CD3xTRP1 markedly increased membranous PD-L1 expression on KPC3-TRP1 cells on all evaluated time points, but was most pronounced at 24 and 72 hours pi.

**Figure 5.**
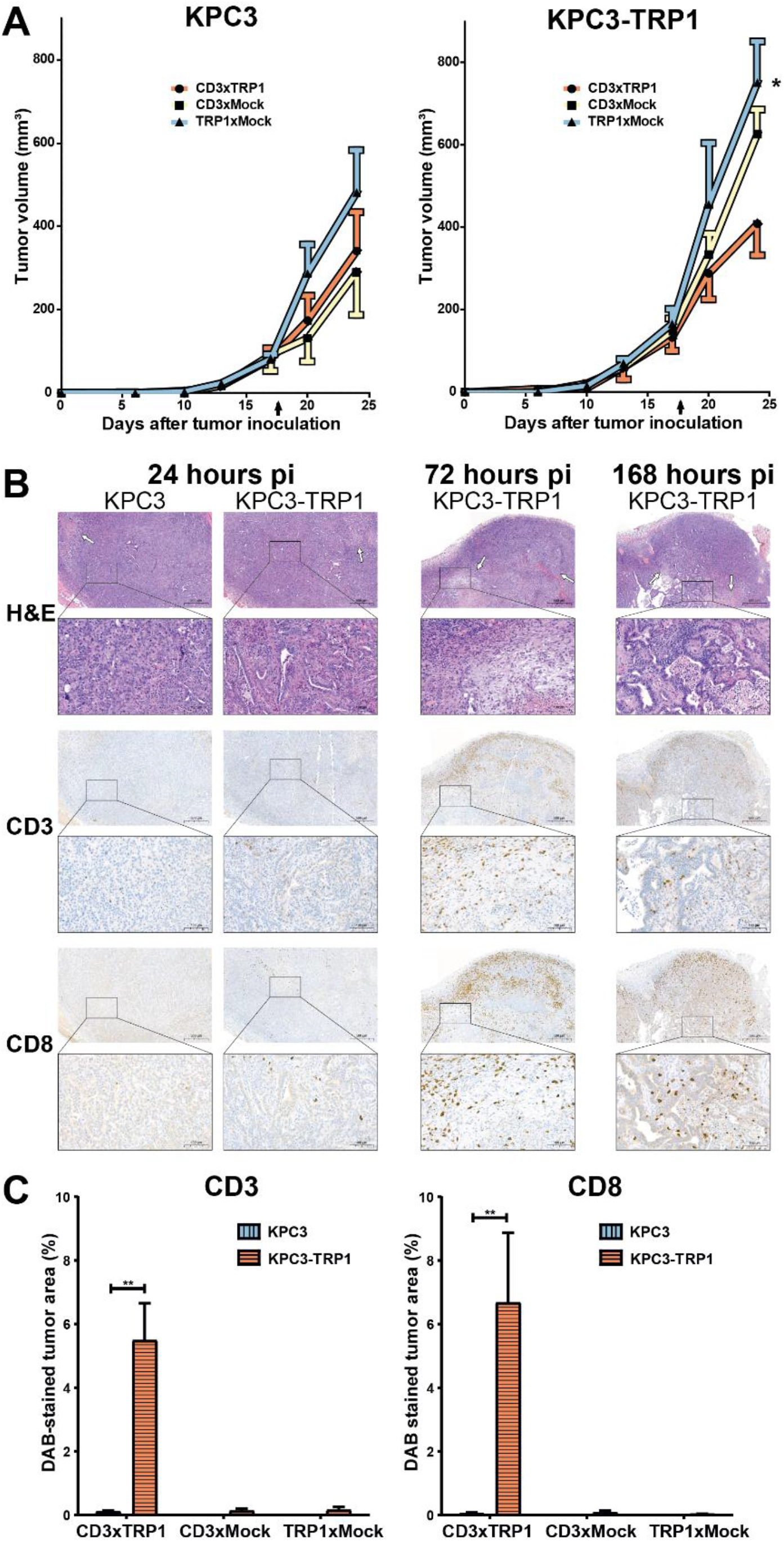
Therapy effects of CD3xTRP1 on KPC3-TRP1 and KPC3 tumors over time. Tumor growth curves and immunohistochemical evaluation of treatment effects in 12.5 µg [^111^In]In/[^125^I]I-CD3xTRP1, [^111^In]In/[^125^I]I-CD3xMock, and [^111^In]In/[^125^I]I-TRP1xMock treated contralateral KPC3-TRP1 and KPC3 tumors-bearing C57BL/6J mice at 24, 72, and 168 hours post intraperitoneal injection. **(A)** Tumor growth curves of KPC3-TRP1 and KPC3 tumors of radiolabeled CD3xTRP1, CD3xMock, and TRP1xMock treated mice of the 168 hour biodistribution group. Arrow indicates time point of bsAb injection. Data shown are mean ± SEM. Differences in KPC3 and KPC3-TRP1^+^ tumor volumes (mean ± SD) were tested for significance using one-way ANOVAs with a Bonferroni *post hoc* test (*=p<0.01). **(B)** Representative images of 5 µm formalin-fixed paraffin-embedded (FFPE) sections of KPC3-TRP1 and KPC3 tumors over time stained for H&E, CD8, CD3, PD-L1, Ly6G, and F4/80. Arrows indicates necrotic tissue regions. The scale bars represents 500 µm, or 100 µm in insert. **(C)** Quantified CD3 and CD8 positively stained areas of KPC3 and KPC3-TRP1 tumors treated with CD3xTRP1, and KPC3-TRP1 treated with CD3xMock, or TRP1xMock, 168 hours pi. Increases in %DAB-stained area were tested for significance using two-tailed t-tests (**=p<0.01).

In CD3xMock treated KPC3-TRP1 tumors, the observed effects were substantially less pronounced, with marginally increased necrotic tumor areas, CD8^+^ T cell numbers, and PD-L1 tumor cell expression only at 24 and 72 hours pi (figure S3). In TRP1xMock treated mice all IHC evaluated features were comparable for KPC3-TRP1 and KPC3 tumors. In none of the KPC3 tumors, the effects observed in KPC3-TRP1 tumors were present. In summary, CD3xTRP1 effectuated anti-tumor immune responses that were predominantly mediated by CD8^+^ lymphocytes and resulted in necrosis, transient upregulation of PD-L1 on tumor cells, and growth delay in KPC3-TRP1^+^ tumors.

## Discussion

Immunotherapeutic CD3-targeting bispecific antibodies cross-link tumor cells with cytotoxic T cells, irrespective of their TCR specificity, thereby inducing local anti-tumor T-cell responses [2]. To further optimize their efficacy, it is important to understand their pharmacokinetics, uptake in tumor and normal tissues, and cellular internalization. Here, we evaluated these aspects *in vivo* using a completely syngeneic mouse model consisting of a full-length radiolabeled murine CD3xTRP1 bsAb, immunocompetent mice which endogenously express the TAA, and murine TRP1-expressing tumors. We demonstrate that CD3xTRP1 distributes to TRP1-positive tumors and CD3-positive tissues, and that a large fraction of CD3xTRP1 is internalized. Nonetheless, anti-tumor immune responses were induced as observed by profound CD8^+^ T-cell influx and delayed tumor growth.

*In vivo* pharmacokinetic SPECT/CT imaging and *ex vivo* biodistribution studies showed that upon intraperitoneal administration, CD3xTRP1 is readily absorbed into the circulation, and distributes specifically towards KPC3-TRP1 tumors and CD3-rich tissues, with uptake peaking 24 hours pi. In a clinically more relevant tumor model B16F10, with endogenous TRP1 expression levels at an order of magnitude lower than KPC3-TRP1 [9], the tumor uptake of CD3xTRP1 was only twice as low, whereas the overall PK and biodistribution was similar. Thus, TAA expression levels influence the absolute tumor-targeted dose, however not necessarily in a one-to-one correlation. Our studies with the control bsAb lacking CD3-affinity, together with the initial absence of tumor-residing CD3^+^ T cells, shows that uptake of CD3xTRP1 in TRP1-positive tumors is primarily TRP1-mediated. Furthermore, circulatory CD3xTRP1 predominantly resided in the serum fraction of blood, indicating that CD3^+^ T cells are generally not traffickers of CD3-bsAbs to the tumor. In pigmented skin, where endogenous TRP1 is expressed by melanocytes, no measurable specific accumulation of CD3xTRP1 was observed. A potential explanation might be that the localization of melanocytes in the avascular epidermis hinders efficient targeting of CD3xTRP1 [20]. This indicates that “on-target, off-tumor” targeting to endogenously expressed TAA does not necessarily affect biodistributions through “antigen sink” effects for this bsAb. Overall, CD3xTRP1’s biodistribution is comparable with other CD3-bsAbs that primarily show uptake in lymphoid organs and TAA-expressing tissues [21, 22] and, as expected, target-tissue uptake is significantly higher compared with molecules with considerably lower circulatory half-lives, such as bispecific T-cell engagers (BiTEs) [23, 24].

SPECT/CT imaging further revealed peritumoral uptake of CD3xTRP1 and control bsAbs around both KPC3 and KPC3-TRP1 but not B16F10 tumors, indicating that it was tumor-type specific and not TRP1- or CD3-mediated. As the Fc-portion of CD3xTRP1 was “LALA” mutated to minimize Fcγ-receptor (FcγR) binding, an FcγR-mediated uptake is also unlikely. However, to definitively exclude FcγRs’ involvement a “PGLALA” mutation could be introduced [25]. Therefore, potential explanations for the non-specific uptake include increased angiogenesis and/or leakiness in the peri-tumoral area, trapping in extracellular matrix deposits induced by tumor cells, or accumulation in extratumoral exudate excreted by the stroma and/or KPC3 cells of pancreatic ductal origin [12, 26].

To exert their therapeutic effect, CD3-bsAbs require simultaneous binding of TAA and CD3 on the membrane of tumor and CD3^+^ T cells, respectively. Therefore, internalization inevitably decreases its therapeutical available dose. Interestingly, for the therapeutically active CD3xTRP1, we observed receptor-mediated internalization in both KPC3-TRP1 tumors and CD3-rich tissues. In the tumor, this decreased the extracellular dose by half within 24 hours and to ∼1/5th within 72 hours pi. In SLOs, even faster internalization resulted in endocytosis of 80% of spleen accumulated CD3xTRP1 within 24 hours. This may be explained by endocytosis signals in CD3ε subunits, which is normally involved in constitutive recycling and downregulating expression of CD3ε, which can be triggered by T-cell activation and antibody binding [27, 28]. Additionally, we observed non-target-mediated internalization of CD3xTRP1, CD3xMock, and TRP1xMock in KPC3 tumors and SLOs, albeit considerably slower. Possible mechanisms include micropinocytosis, which occurs in T cells and pancreatic ductal adenocarcinomas [29, 30], and phagocytosis of tumor or immune-cell debris by tumor-infiltrated macrophages. Others have also observed internalization of CD3-bsAbs in tumors and SLOs at a fixed 72 hours pi time point using a CD3-bsAb targeting human-CD3ε in transgenic mice expressing both mouse and human CD3ε [21]. We expand upon that knowledge by providing a longitudinal overview of this process with our fully functional mouse CD3-bsAb in a non-artificial setting. Previously, upon administering CD3xTRP1, transient T-cell activation was observed throughout the body, except for those in the tumor, which stayed activated for at least 4 days [9]. This might be explained by the higher internalization rate in SLOs compared to KPC3-TRP1 tumors, which suggests that balancing CD3-bsAb internalization rates can help mitigate immune related adverse effects (IRAEs), while still inducing prolonged activation of anti-tumor T-cell responses.

Despite rapidly internalizing, CD3xTRP1 managed to elicit anti-tumor immune responses as KPC3-TRP1 tumors showed delayed growth, influx of CD8^+^ T cells, increases in necrotic areas, and a subsequent influx of neutrophils. These results correspond with previous findings for CD3xTRP1 [9]. Additionally, we observed CD3xTRP1-mediated upregulation of the immune checkpoint PD-L1 on TRP1-positive tumors, which is probably an effect of interferons produced by activated immune cells. This corresponds with previous (pre)clinical observations and thus supports the proposal of combining CD3-bsAbs with immune checkpoint inhibitors to increase anti-tumor efficacy [31-33].

Although CD3xTRP1 shows favorable distribution characteristics resulting in effective anti-tumor immune responses, this study does show opportunities for improving CD3-bsAb immunotherapy by 1) increasing tumor uptake, and 2) reducing internalization rates. To increase absolute CD3xTRP1 tumor uptake, shifting uptake from SLOs toward the tumor is preferred above increasing the administered dose as both the tumor and SLOs uptake were not saturated. For high TAA-expressing tumors (e.g. KPC3-TRP1) this can be accomplished by decreasing the CD3-affinity of the bsAb, whereas for low TAA expressing-tumors the resulting decrease in therapeutic potency can prohibit this approach [22, 33]. Additionally, lowering CD3-affinities may reduce CD3-bsAbs’ IRAE potential [22, 34]. Other methods to increase tumor uptake include using engineered antibodies formats, such as asymmetric “2-to-1” bsAbs, or using masked CD3-binding arms which are unmasked within the tumor to prevent “on-target, off-tumor” binding [35]. This can also limit activation-induced cell death in SLOs through Abs:CD3-binding, thereby increasing the number of healthy non-anergic T cells that can be recruited for anti-tumor responses [36].

A second approach to improve CD3-bsAb immunotherapy is through reducing internalization rates. In the tumor, reduced internalization increases the extracellular available dose, thereby mitigating the loss of therapeutic efficacy. This may be accomplished by reducing the TAA-affinity, as it has been shown that internalization rate increases with epitope affinity for other targets [37]. In SLOs, when not subsequently retained through subsite rebinding, reduced internalization could result in an increase in CD3-bsAb in circulation, which consequently becomes available for tumor targeting. Given the many different CD3-bsAb formats that have been developed, which all have different TAA targets, PKs, biodistribution, internalization characteristics, and dosing schedules, it is difficult to provide a clear generalization of our findings to these different formats. [21, 22, 38]. Nevertheless, we highly recommend using nuclear imaging for clinical CD3-bsAb candidates to obtain insights into their distribution to tumors and healthy tissues, optimal treatment doses and dosing schedules.

In summary, we used molecular imaging to assess the pharmacokinetics and biodistribution of CD3xTRP1 bsAb and demonstrate specific and efficient accumulation in both transfected and endogenously TRP1-expressing tumors and CD3-rich tissues in fully syngeneic immunocompetent mice. We further demonstrate that CD3^+^ T-cell dominated anti-tumor immune responses can be achieved despite rapid CD3xTRP1 internalization. We are intrigued whether reducing internalization of CD3-bsAbs and/or shifting their uptake towards the tumor can further increase their therapeutic efficacy.

## Declarations

## Acknowledgments

We thank Vitalijs Ovcinnikovs, Katy Lloyd, Janine Schuurman, Renoud Marijnissen, and Kristel Kemper (all Genmab BV employees) for kindly providing the CD3-bsAbs and valuable discussions. We thank Milou Boswinkel, Annemarie Kip, Cathelijne Frielink, Bianca Lemmers-van de Weem, Iris Lamers-Elemans, Floor Moonen, and Kitty Lemmens-Hermans for technical assistance with the animal experiments, and Simone Kleinendorst for co-creating the IHC analysis macro.

## Authors’ contributions

Conceptualization: GGWS, JM, TvH and SH. Methodology: GGWS, JM, EA, JMK, TvH and SH. Software: GGWS. Validation: GGWS, JM. Formal analysis: GGWS, JM, JMK, EW. Investigation: GGWS, JM, EW, and JMK. Resources: JM, TvH and SH. Data Curation: GGWS, JM, EW, and JMK. Writing – original draft preparation: GGWS. Writing – review and editing: GGWS, EW, JMK, JM, EA, TvH, and SH. Visualization: GGWS, JM. Supervision: EA, TvH and SH. Project administration: GGWS, JM. Funding acquisition: TvH and SH. Guarantor: SH.

## Funding

This research received funding from the Dutch Research Council (SH, NWO; 09150172010054) and from Genmab via a research grant (TvH).

## Competing interest

TvH received a research grant from Genmab. Bispecific antibodies were provided by Genmab. TvH and JM are authors on a patent involving the combination of CD3-bsAb therapy in combination with vaccination. The other authors declare that they have no competing interests.

## Ethics approval

All *in vivo* experiments were approved by the Animal Welfare Body of the Radboud University, Nijmegen, and the Central Authority for Scientific Procedures on Animals (AVD1030020209645) and were performed in accordance with the principles stated by the Dutch Act on Animal Experiments (2014).

### Abbreviations

CD3-bsAbs: CD3 bispecific antibodies
TAA: tumor-associated antigen
TRP1: tyrosinase related protein 1
PK: pharmacokinetics
SLO: secondary lymphoid organ
FCS: fetal calf serum
ITC-DTPA: isothiocyanatobenzyl-diethylenetriaminepentaacetic acid
RT: room temperature
^111^In: indium-111
^125^I: iodine-125
(M)Bq: (Mega)bequerel
EDTA: ethylenediaminetetraacetic
iTLC: instant thin-layer chromatography
PBS: phosphate buffered saline
LDH: lactate dehydrogenase
BSA: bovine serum albumin
pi: post injection
%ID/g: percentage injected dose per gram
SPECT: single-photon emission computed tomography
MIP: maximum-intensity projections
BiTEs: bispecific T-cell engagers
FcγR: Fcγ-receptor
IRAEs: immune related adverse effects
FFPE: Formalin-fixed paraffin-embedded
PD-L1: Programmed death-ligand 1

