## Supplemental figures and tables for "Imaging the pharmacokinetics and therapeutic availability of the bispecific CD3xTRP1 antibody in syngeneic mouse tumor models"

### Supplementals

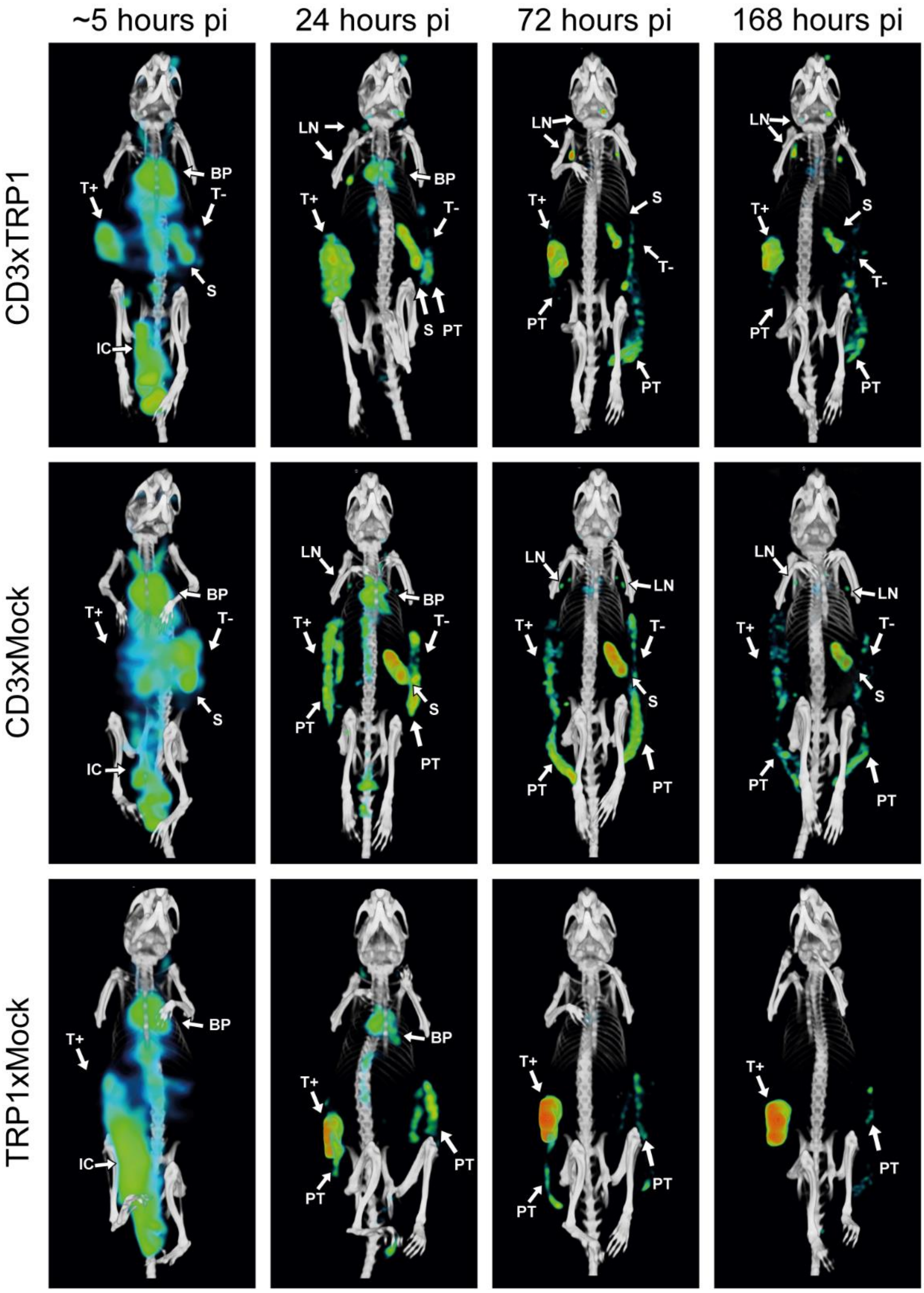


**Figure S1. SPECT/CT images showing the *in vivo* biodistribution of CD3xTRP1, CD3xMock, and TRP1xMock over time.** Representative MIPs of the *in vivo* biodistribution of 12.5 µg **(A)** [$^{111}$In]In–CD3xTRP1, **(B)** [$^{111}$In]In–CD3xMock, and **(C)** [$^{111}$In]In–TRP1xMock in C57BL/6J mice bearing KPC3 and KPC3-TRP1 tumors on contralateral flanks at ~5, 24, 72, and 168 hours post intraperitoneal injection. Tissue uptake of [$^{111}$In]In–bsAbs is visualized with MIPs generated from microSPECT/CT data using the same thresholds at all time point. Indicated tissues with [$^{111}$In]In-bsAb uptake are: blood pool (BP), intraperitoneal cavity (IC), KPC3-TRP1 tumor (T+), KPC3 tumor (T-), spleen (S), lymph nodes (LN), and peri- and extratumoral accumulation (PT).

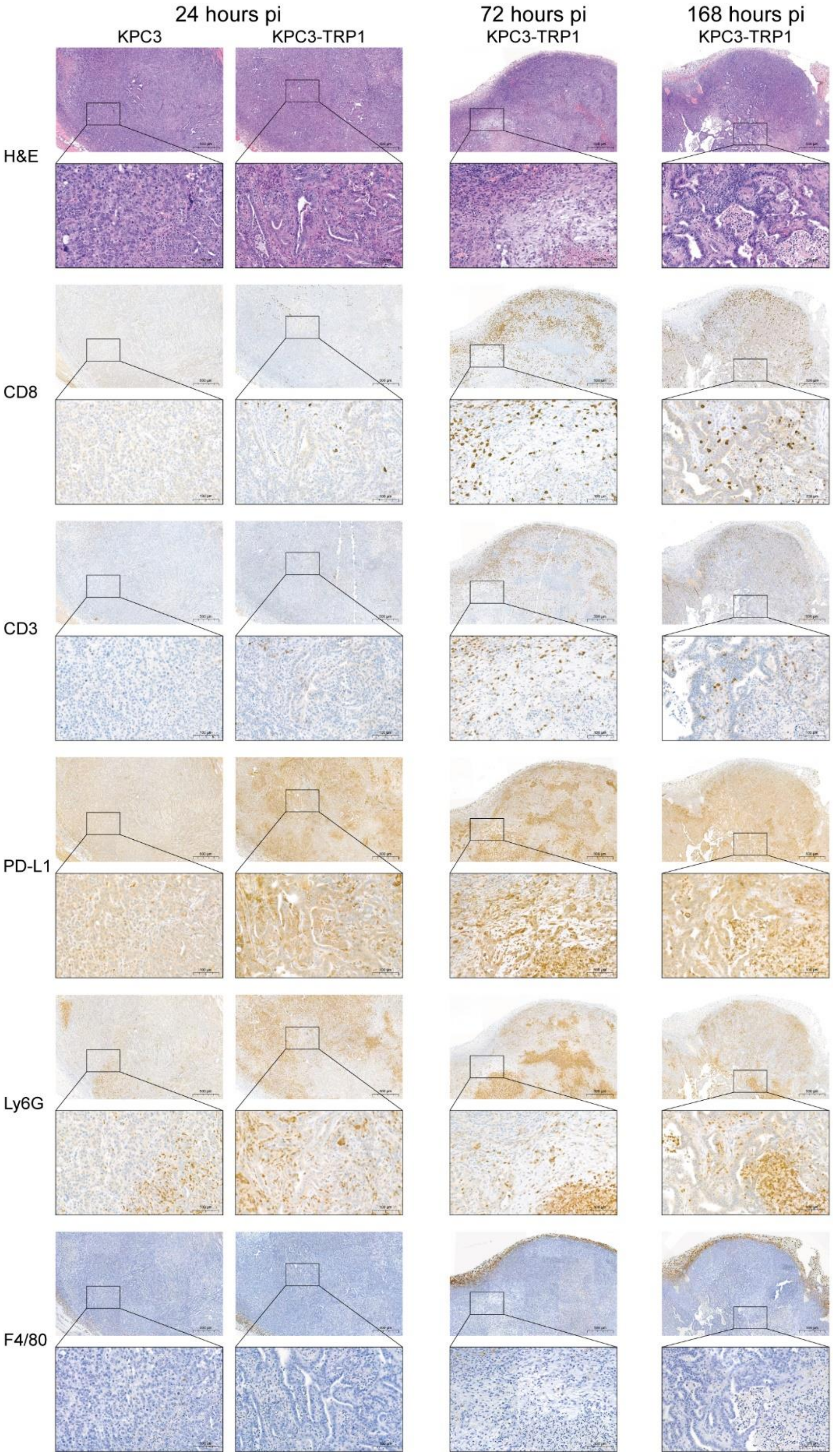


**Figure S2. Therapy effects of CD3xTRP1 on KPC3-TRP1 and KPC3 tumors over time.**
Immunohistochemistry on 5 µm sections of FFPE KPC3-TRP1 and KPC3 tumors of contralaterally tumor-bearing C57BL/6J mice treated with 12.5 µg [$^{111}$In]In/[$^{125}$I]I-CD3xTRP1 at 24, 72, and 168 hours post intraperitoneal injection. Representative images of H&E, CD8, CD3, PD-L1, Ly6G, and F4/80 stains. The scale bars represents 500 µm, or 100 µm in insert.

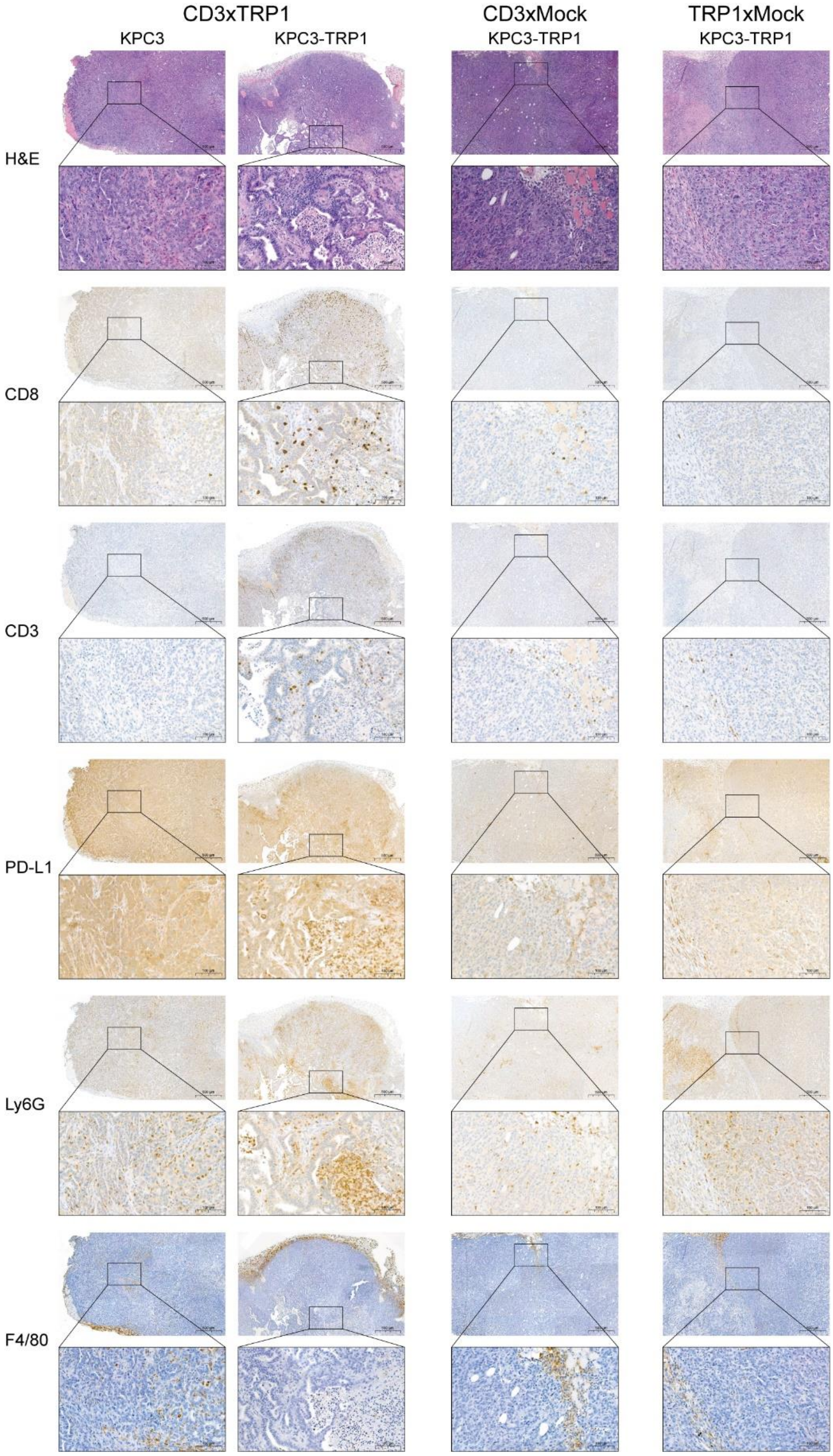


**Figure S3. Therapy effects of CD3xTRP1, CD3xMock, and TRP1xMock on KPC3-TRP1 tumors.** Immunohistochemistry on 5 µm sections of FFPE KPC3 and KPC3-TRP1 tumors of contralaterally tumor-bearing C57BL/6J mice treated with 12.5 µg [$^{111}$In]In/[$^{125}$I]I-CD3xTRP1, [$^{111}$In]In/[$^{125}$I]I-CD3xMock, or [$^{111}$In]In/[$^{125}$I]I-TRP1xMock at 168 hours post intraperitoneal injection. Representative images of H&E, CD8, CD3, PD-L1, Ly6G, and F4/80 stains. The scale bars represents 500 µm, or 100 µm in insert.

**Table S1.** *Ex vivo* biodistribution analysis of [$^{111}$In]In-CD3xTRP1, [$^{111}$In]In-CD3xMock, and [$^{111}$In]In-TRP1xMock in KPC3-TRP1 and KPC3 tumor-bearing C57BL/6J mice at 24, 72, and 168 hours pi, and [$^{111}$In]In-CD3xTRP1 in B16F10 tumor-bearing C57BL/6J mice. Mean results, standard deviation, and sample size per tissue are shown in %ID/g (mean ± sd (n=x)).

| **24 hours pi** | **[$^{111}$In]In-CD3xTRP1** | **[$^{111}$In]In-CD3xMock** | **[$^{111}$In]In-TRP1xMock** |
|---|---|---|---|
| | (%ID/g) | (%ID/g) | (%ID/g) |
| **Blood** | 22.3 ± 2.1 (n=5) | 28.6 ± 7.8 (n=7) | 32.1 ± 4.7 (n=6) |
| **Skin (pigmented)** | 18.3 ± 0.0 (n=1) | 11.5 ± 6.0 (n=5) | 13.7 ± 1.3 (n=3) |
| **Skin (non-pigmented)** | 9.6 ± 4.5 (n=5) | 11.3 ± 4.8 (n=7) | 15.7 ± 12.3 (n=6) |
| **Brown adipose tissue** | 3.4 ± 0.5 (n=5) | 4.3 ± 1.5 (n=7) | 5.5 ± 1.6 (n=6) |
| **Lymph nodes** | 50.3 ± 14.8 (n=5) | 50.1 ± 19.2 (n=7) | 19.2 ± 15.2 (n=6) |
| **KPC3** | 8.3 ± 1.3 (n=4) | 13.0 ± 8.4 (n=7) | 10.6 ± 2.2 (n=5) |
| **KPC3-TRP1** | 37.7 ± 5.3 (n=5) | 10.0 ± 3.2 (n=6) | 54.8 ± 15.3 (n=6) |
| **Muscle** | 1.3 ± 0.2 (n=5) | 1.5 ± 0.4 (n=7) | 2.0 ± 0.5 (n=6) |
| **Thymus** | 8.0 ± 1.0 (n=5) | 10.8 ± 2.3 (n=7) | 7.1 ± 2.1 (n=6) |
| **Heart** | 5.3 ± 0.5 (n=5) | 6.2 ± 2.2 (n=7) | 7.8 ± 1.9 (n=6) |
| **Lung** | 10.9 ± 1.9 (n=5) | 13.4 ± 3.5 (n=7) | 15.4 ± 5.0 (n=6) |
| **Spleen** | 29.0 ± 3.9 (n=5) | 48.7 ± 14.7 (n=7) | 5.1 ± 1.1 (n=6) |
| **Pancreas** | 4.8 ± 0.7 (n=5) | 5.7 ± 1.5 (n=7) | 6.8 ± 1.5 (n=6) |
| **Stomach** | 3.2 ± 0.5 (n=5) | 4.0 ± 1.1 (n=7) | 4.9 ± 1.0 (n=6) |
| **Duodenum** | 5.1 ± 0.8 (n=5) | 7.0 ± 2.0 (n=7) | 5.9 ± 1.3 (n=6) |
| **Colon** | 2.7 ± 0.2 (n=5) | 3.8 ± 1.3 (n=7) | 4.0 ± 0.6 (n=6) |
| **Liver** | 3.3 ± 0.5 (n=5) | 4.3 ± 0.9 (n=7) | 4.4 ± 0.6 (n=6) |
| **Kidney** | 8.9 ± 0.8 (n=5) | 10.4 ± 2.8 (n=7) | 12.1 ± 2.0 (n=6) |
| **Prostate** | 5.9 ± 1.6 (n=5) | 8.5 ± 3.4 (n=7) | 9.0 ± 2.8 (n=6) |
| **Bone marrow** | 9.9 ± 2.6 (n=5) | 12.8 ± 4.0 (n=7) | 11.3 ± 3.3 (n=6) |
| **Bone** | 2.2 ± 0.1 (n=5) | 3.1 ± 0.8 (n=7) | 2.8 ± 0.7 (n=6) |

| | KPC3-TRP1 and KPC | | | B16F10 |
|---|---|---|---|---|
| **72 hours pi** | **[$^{111}$In]In-CD3xTRP1** | **[$^{111}$In]In-CD3xMock** | **[$^{111}$In]In-TRP1xMock** | **[$^{111}$In]In-CD3xTRP1** |
| | (%ID/g) | (%ID/g) | (%ID/g) | (%ID/g) |
| **Blood** | 16.3 ± 5.4 (n=5) | 19.9 ± 4.9 (n=7) | 23.8 ± 3.0 (n=6) | 17.6 ± 2.3 (n=4) |
| **Skin (pigmented)** | 9.0 ± 1.9 (n=2) | 9.4 ± 2.5 (n=5) | 12.9 ± 1.8 (n=2) | 7.7 ± 2.2 (n=4) |
| **Skin (non-pigmented)** | 8.3 ± 3.0 (n=5) | 10.6 ± 4.8 (n=7) | 15.5 ± 4.2 (n=6) | 8.6 ± 5.1 (n=4) |
| **Brown adipose tissue** | 2.7 ± 0.9 (n=5) | 3.8 ± 1.0 (n=7) | 4.0 ± 0.5 (n=6) | 2.7 ± 0.5 (n=4) |
| **Lymph nodes** | 45.3 ± 13.6 (n=5) | 63.2 ± 17.2 (n=7) | 11.5 ± 1.4 (n=6) | 30.2 ± 11.3 (n=4) |
| **KPC3** | 6.2 ± 2.1 (n=5) | 9.1 ± 2.7 (n=7) | 11.2 ± 1.4 (n=6) | n/a |
| **KPC3-TRP1 or B16F10** | 33.5 ± 15.4 (n=5) | 11.5 ± 7.3 (n=7) | 74.2 ± 15.4 (n=6) | 25.1 ± 15.1 (n=4) |
| **Muscle** | 1.0 ± 0.3 (n=5) | 1.2 ± 0.3 (n=7) | 1.8 ± 0.3 (n=6) | 0.7 ± 0.1 (n=4) |
| **Thymus** | 10.0 ± 3.1 (n=5) | 13.4 ± 3.4 (n=7) | 5.2 ± 1.0 (n=6) | 10.9 ± 2.1 (n=4) |
| **Heart** | 3.6 ± 1.2 (n=5) | 4.4 ± 1.1 (n=7) | 5.0 ± 0.9 (n=6) | 4.1 ± 0.9 (n=4) |
| **Lung** | 8.7 ± 3.9 (n=5) | 10.9 ± 2.7 (n=7) | 11.4 ± 2.5 (n=6) | 7.4 ± 1.3 (n=4) |
| **Spleen** | 23.8 ± 10.4 (n=5) | 46.4 ± 17.7 (n=7) | 5.5 ± 0.5 (n=6) | 30.9 ± 8.2 (n=4) |
| **Pancreas** | 2.7 ± 0.9 (n=5) | 3.3 ± 0.6 (n=7) | 3.7 ± 0.5 (n=6) | 2.6 ± 0.3 (n=4) |
| **Stomach** | 2.3 ± 0.9 (n=5) | 2.8 ± 0.6 (n=7) | 3.0 ± 0.8 (n=6) | 1.8 ± 0.5 (n=4) |
| **Duodenum** | 4.4 ± 1.8 (n=5) | 5.5 ± 1.4 (n=7) | 3.3 ± 0.7 (n=6) | 3.8 ± 0.5 (n=4) |
| **Colon** | 1.7 ± 0.5 (n=5) | 2.4 ± 0.6 (n=7) | 2.2 ± 0.5 (n=6) | 1.7 ± 0.3 (n=4) |
| **Liver** | 3.8 ± 1.2 (n=5) | 4.4 ± 1.0 (n=7) | 3.9 ± 0.3 (n=6) | 5.1 ± 0.7 (n=4) |
| **Kidney** | 6.5 ± 2.0 (n=5) | 8.2 ± 1.8 (n=7) | 9.9 ± 1.1 (n=6) | 6.6 ± 0.7 (n=4) |
| **Prostate** | 3.5 ± 1.5 (n=5) | 4.2 ± 1.4 (n=7) | 5.4 ± 1.0 (n=6) | 3.0 ± 0.6 (n=4) |
| **Bone marrow** | 5.3 ± 1.7 (n=5) | 8.7 ± 2.4 (n=7) | 6.8 ± 1.0 (n=6) | 4.8 ± 0.8 (n=4) |
| **Bone** | 1.7 ± 0.6 (n=5) | 2.6 ± 0.8 (n=7) | 2.6 ± 0.5 (n=6) | 1.5 ± 0.2 (n=4) |

| 168 hours pi | [$^{111}$In]In-CD3xTRP1 | [$^{111}$In]In-CD3xMock | [$^{111}$In]In-TRP1xMock |
|---|---|---|---|
| | (%ID/g) | (%ID/g) | (%ID/g) |
| **Blood** | 7.8 ± 3.7 (n=6) | 11.9 ± 3.5 (n=5) | 9.4 ± 2.6 (n=6) |
| **Skin (pigmented)** | 3.8 ± 2.1 (n=5) | 4.0 ± 1.6 (n=3) | 6.4 ± 1.2 (n=6) |
| **Skin (non-pigmented)** | 4.9 ± 2.9 (n=6) | 6.1 ± 3.3 (n=5) | 8.5 ± 2.5 (n=7) |
| **Brown adipose tissue** | 1.6 ± 0.7 (n=6) | 2.8 ± 1.0 (n=5) | 2.1 ± 0.6 (n=7) |
| **Lymph nodes** | 26.3 ± 10.7 (n=6) | 48.9 ± 30.3 (n=5) | 9.7 ± 2.2 (n=7) |
| **KPC3** | 3.3 ± 1.5 (n=6) | 5.2 ± 1.9 (n=4) | 4.2 ± 0.8 (n=7) |
| **KPC3-TRP1** | 15.1 ± 5.8 (n=6) | 5.2 ± 1.7 (n=5) | 32.0 ± 10.1 (n=7) |
| **Muscle** | 0.6 ± 0.3 (n=6) | 0.7 ± 0.3 (n=5) | 0.7 ± 0.1 (n=7) |
| **Thymus** | 7.7 ± 3.0 (n=6) | 11.5 ± 3.6 (n=5) | 2.5 ± 0.4 (n=7) |
| **Heart** | 2.0 ± 0.9 (n=6) | 2.7 ± 0.9 (n=5) | 2.3 ± 0.6 (n=7) |
| **Lung** | 4.0 ± 2.0 (n=6) | 5.9 ± 2.0 (n=5) | 4.3 ± 0.6 (n=7) |
| **Spleen** | 13.9 ± 7.0 (n=6) | 27.6 ± 11.0 (n=5) | 2.8 ± 0.5 (n=7) |
| **Pancreas** | 1.7 ± 0.7 (n=6) | 2.3 ± 0.9 (n=5) | 1.7 ± 0.4 (n=7) |
| **Stomach** | 1.1 ± 0.5 (n=6) | 1.8 ± 0.7 (n=5) | 1.3 ± 0.4 (n=7) |
| **Duodenum** | 2.5 ± 1.1 (n=6) | 4.6 ± 1.4 (n=5) | 2.0 ± 0.6 (n=7) |
| **Colon** | 0.9 ± 0.4 (n=6) | 1.4 ± 0.5 (n=5) | 1.1 ± 0.3 (n=7) |
| **Liver** | 2.4 ± 1.0 (n=6) | 3.4 ± 1.0 (n=5) | 2.2 ± 0.3 (n=7) |
| **Kidney** | 3.8 ± 1.5 (n=6) | 5.2 ± 1.4 (n=5) | 5.1 ± 0.9 (n=7) |
| **Prostate** | 1.7 ± 0.7 (n=6) | 2.6 ± 1.0 (n=5) | 2.2 ± 0.4 (n=7) |
| **Bone marrow** | 2.7 ± 1.1 (n=6) | 5.0 ± 1.7 (n=5) | 2.7 ± 0.6 (n=7) |
| **Bone** | 0.9 ± 0.3 (n=6) | 1.6 ± 0.5 (n=5) | 1.1 ± 0.2 (n=7) |

**Table S2.** Absolute accumulated CD3xTRP1 as observed with *ex vivo* biodistribution of [$^{111}$In]In-CD3xTRP1 in KPC3-TRP1 and KPC3 tumor-bearing C57BL/6J mice at 24, 72, and 168 hours pi. The %ID which is not corrected for tissue weight is shown only for those tissues that were excised at a whole. Mean results, standard deviation, and sample size per tissue are shown in %ID (mean ± sd (n=x)).

| | **24 hours pi** | **72 hours pi** | **168 hours pi** |
|---|---|---|---|
| | (%ID/g) | (%ID/g) | (%ID/g) |
| **KPC3** | 0.5 ± 0.4 (n=5) | 1.6 ± 1.1 (n=5) | 1.2 ± 1.0 (n=6) |
| **KPC3-TRP1** | 5.9 ± 2.2 (n=5) | 6.5 ± 1.9 (n=5) | 6.0 ± 2.9 (n=6) |
| **Spleen** | 2.7 ± 0.3 (n=5) | 2.6 ± 0.8 (n=5) | 2.1 ± 0.9 (n=6) |
| **Lymph nodes** | 0.2 ± 0.0 (n=5) | 0.3 ± 0.1 (n=5) | 0.3 ± 0.1 (n=6) |

**Table S3.** *Ex vivo* biodistribution analysis of $^{125}$I-labeled CD3xTRP1, CD3xMock, and TRP1xMock in KPC3-TRP1 and KPC3 tumor-bearing C57BL/6J mice at 24, 72, and 168 hours pi. Mean results, standard deviation, and sample size per tissue are shown in %ID/g (mean ± sd (n=x)).

| **24 hours pi** | **[$^{125}$I]I-CD3xTRP1** | **[$^{125}$I]I-CD3xMock** | **[$^{125}$I]I-TRP1xMock** |
|---|---|---|---|
| | (%ID/g) | (%ID/g) | (%ID/g) |
| **Blood** | 18.5 ± 2.1 (n=5) | 22.5 ± 6.3 (n=7) | 21.6 ± 3.2 (n=6) |
| **Skin (pigmented)** | 14.7 ± 0.0 (n=1) | 8.3 ± 3.9 (n=5) | 8.9 ± 0.6 (n=3) |
| **Skin (non-pigmented)** | 7.6 ± 3.6 (n=5) | 8.3 ± 3.4 (n=7) | 9.4 ± 6.4 (n=6) |
| **Brown adipose tissue** | 2.8 ± 0.4 (n=5) | 3.3 ± 1.1 (n=7) | 3.6 ± 0.9 (n=6) |
| **Lymph nodes** | 12.8 ± 2.2 (n=5) | 12.0 ± 3.2 (n=7) | 7.5 ± 5.2 (n=6) |
| **KPC3** | 5.9 ± 0.9 (n=4) | 8.6 ± 6.0 (n=7) | 5.9 ± 1.2 (n=5) |
| **KPC3-TRP1** | 17.0 ± 2.7 (n=5) | 6.6 ± 2.0 (n=6) | 23.3 ± 6.2 (n=6) |
| **Muscle** | 1.1 ± 0.2 (n=5) | 1.2 ± 0.4 (n=7) | 1.3 ± 0.3 (n=6) |
| **Thymus** | 3.7 ± 0.8 (n=5) | 4.8 ± 1.2 (n=7) | 4.5 ± 1.4 (n=6) |
| **Heart** | 4.3 ± 0.4 (n=5) | 4.9 ± 1.8 (n=7) | 5.2 ± 1.4 (n=6) |
| **Lung** | 8.8 ± 1.7 (n=5) | 10.0 ± 2.8 (n=7) | 10.0 ± 3.3 (n=6) |
| **Spleen** | 5.9 ± 0.6 (n=5) | 7.8 ± 2.4 (n=7) | 2.7 ± 0.6 (n=6) |
| **Pancreas** | 3.2 ± 0.4 (n=5) | 3.8 ± 1.0 (n=7) | 4.0 ± 0.9 (n=6) |
| **Stomach** | 2.7 ± 1.0 (n=5) | 2.9 ± 1.0 (n=7) | 3.5 ± 0.6 (n=6) |
| **Duodenum** | 4.7 ± 2.9 (n=5) | 5.7 ± 3.9 (n=7) | 3.8 ± 0.9 (n=6) |
| **Colon** | 1.9 ± 0.2 (n=5) | 2.6 ± 1.0 (n=7) | 2.4 ± 0.4 (n=6) |
| **Liver** | 2.1 ± 0.3 (n=5) | 2.6 ± 0.6 (n=7) | 2.6 ± 0.4 (n=6) |
| **Kidney** | 5.3 ± 0.4 (n=5) | 6.3 ± 1.8 (n=7) | 6.1 ± 1.0 (n=6) |
| **Prostate** | 5.1 ± 1.7 (n=5) | 6.5 ± 2.5 (n=7) | 5.9 ± 1.9 (n=6) |
| **Bone marrow** | 5.8 ± 1.2 (n=5) | 6.4 ± 2.0 (n=7) | 6.5 ± 2.2 (n=6) |
| **Bone** | 1.4 ± 0.1 (n=5) | 1.8 ± 0.5 (n=7) | 1.6 ± 0.4 (n=6) |

| 72 hours pi | [$^{125}$I]I-CD3xTRP1 | [$^{125}$I]I-CD3xMock | [$^{125}$I]I-TRP1xMock |
|---|---|---|---|
| | (%ID/g) | (%ID/g) | (%ID/g) |
| **Blood** | 13.7 ± 4.5 (n=5) | 16.1 ± 3.9 (n=7) | 15.9 ± 2.1 (n=6) |
| **Skin (pigmented)** | 6.1 ± 1.2 (n=2) | 6.9 ± 1.6 (n=5) | 7.5 ± 0.8 (n=2) |
| **Skin (non-pigmented)** | 5.9 ± 2.4 (n=5) | 8.0 ± 3.5 (n=7) | 9.2 ± 2.5 (n=6) |
| **Brown adipose tissue** | 2.2 ± 0.8 (n=5) | 2.8 ± 0.7 (n=7) | 2.5 ± 0.3 (n=6) |
| **Lymph nodes** | 6.8 ± 2.0 (n=5) | 8.1 ± 2.3 (n=7) | 3.3 ± 0.3 (n=6) |
| **KPC3** | 3.8 ± 1.4 (n=5) | 5.5 ± 1.6 (n=7) | 5.6 ± 0.9 (n=6) |
| **KPC3-TRP1** | 7.6 ± 4.1 (n=5) | 5.4 ± 1.6 (n=7) | 19.9 ± 3.6 (n=6) |
| **Muscle** | 0.9 ± 0.3 (n=5) | 0.9 ± 0.2 (n=7) | 1.2 ± 0.2 (n=6) |
| **Thymus** | 2.5 ± 0.8 (n=5) | 3.1 ± 0.7 (n=7) | 2.8 ± 0.6 (n=6) |
| **Heart** | 2.9 ± 1.0 (n=5) | 3.4 ± 0.8 (n=7) | 3.3 ± 0.6 (n=6) |
| **Lung** | 6.7 ± 3.0 (n=5) | 8.0 ± 2.0 (n=7) | 7.3 ± 1.7 (n=6) |
| **Spleen** | 3.2 ± 1.3 (n=5) | 4.3 ± 1.2 (n=7) | 2.1 ± 0.2 (n=6) |
| **Pancreas** | 1.7 ± 0.4 (n=5) | 2.0 ± 0.3 (n=7) | 1.9 ± 0.3 (n=6) |
| **Stomach** | 1.9 ± 0.7 (n=5) | 2.2 ± 0.4 (n=7) | 2.2 ± 0.5 (n=6) |
| **Duodenum** | 2.1 ± 0.7 (n=5) | 2.3 ± 0.5 (n=7) | 1.9 ± 0.4 (n=6) |
| **Colon** | 1.2 ± 0.4 (n=5) | 1.4 ± 0.2 (n=7) | 1.3 ± 0.3 (n=6) |
| **Liver** | 1.9 ± 0.6 (n=5) | 2.4 ± 0.8 (n=7) | 2.0 ± 0.1 (n=6) |
| **Kidney** | 3.9 ± 1.1 (n=5) | 4.5 ± 1.4 (n=7) | 4.7 ± 0.6 (n=6) |
| **Prostate** | 2.5 ± 1.0 (n=5) | 3.1 ± 0.8 (n=7) | 3.2 ± 0.6 (n=6) |
| **Bone marrow** | 2.6 ± 0.9 (n=5) | 3.8 ± 0.9 (n=7) | 3.4 ± 0.5 (n=6) |
| **Bone** | 1.0 ± 0.3 (n=5) | 1.3 ± 0.4 (n=7) | 1.3 ± 0.2 (n=6) |

| 168 hours pi | [125I]I-CD3xTRP1 | [125I]I-CD3xMock | [125I]I-TRP1xMock |
|---|---|---|---|
| | (%ID/g) | (%ID/g) | (%ID/g) |
| **Blood** | 6.5 ± 3.1 (n=6) | 9.3 ± 2.6 (n=5) | 6.0 ± 1.7 (n=6) |
| **Skin (pigmented)** | 2.5 ± 1.3 (n=5) | 2.7 ± 1.1 (n=3) | 3.3 ± 0.6 (n=6) |
| **Skin (non-pigmented)** | 3.1 ± 1.7 (n=6) | 4.1 ± 2.3 (n=5) | 4.3 ± 1.3 (n=7) |
| **Brown adipose tissue** | 1.2 ± 0.5 (n=6) | 1.9 ± 0.8 (n=5) | 1.2 ± 0.4 (n=7) |
| **Lymph nodes** | 2.5 ± 1.1 (n=6) | 4.7 ± 2.8 (n=5) | 1.3 ± 0.2 (n=7) |
| **KPC3** | 1.9 ± 0.9 (n=6) | 2.6 ± 0.9 (n=4) | 1.7 ± 0.3 (n=7) |
| **KPC3-TRP1** | 3.1 ± 1.1 (n=6) | 2.8 ± 0.9 (n=5) | 6.2 ± 1.9 (n=7) |
| **Muscle** | 0.4 ± 0.2 (n=6) | 0.5 ± 0.2 (n=5) | 0.4 ± 0.1 (n=7) |
| **Thymus** | 1.4 ± 0.7 (n=6) | 1.7 ± 0.3 (n=5) | 0.9 ± 0.2 (n=7) |
| **Heart** | 1.6 ± 0.8 (n=6) | 2.0 ± 0.6 (n=5) | 1.4 ± 0.4 (n=7) |
| **Lung** | 2.9 ± 1.6 (n=6) | 3.8 ± 1.3 (n=5) | 2.3 ± 0.3 (n=7) |
| **Spleen** | 1.3 ± 0.7 (n=6) | 1.9 ± 0.8 (n=5) | 0.6 ± 0.1 (n=7) |
| **Pancreas** | 0.8 ± 0.4 (n=6) | 1.2 ± 0.4 (n=5) | 0.8 ± 0.2 (n=7) |
| **Stomach** | 0.9 ± 0.3 (n=6) | 1.6 ± 0.4 (n=5) | 1.1 ± 0.2 (n=7) |
| **Duodenum** | 1.0 ± 0.4 (n=6) | 1.4 ± 0.4 (n=5) | 1.0 ± 0.3 (n=7) |
| **Colon** | 0.5 ± 0.3 (n=6) | 0.8 ± 0.3 (n=5) | 0.5 ± 0.2 (n=7) |
| **Liver** | 1.0 ± 0.5 (n=6) | 1.1 ± 0.4 (n=5) | 0.8 ± 0.2 (n=7) |
| **Kidney** | 2.2 ± 0.8 (n=6) | 2.9 ± 0.8 (n=5) | 2.0 ± 0.5 (n=7) |
| **Prostate** | 1.3 ± 0.6 (n=6) | 1.7 ± 0.7 (n=5) | 1.2 ± 0.1 (n=7) |
| **Bone marrow** | 0.9 ± 0.4 (n=6) | 1.4 ± 0.6 (n=5) | 0.8 ± 0.1 (n=7) |
| **Bone** | 0.4 ± 0.1 (n=6) | 0.6 ± 0.2 (n=5) | 0.4 ± 0.1 (n=7) |

**Table S4.** *In vivo* internalization shown as a ratio between $^{125}$I-labeled and $^{111}$In-labeled CD3xTRP1, CD3xMock, and TRP1xMock in KPC3-TRP1 and KPC3 tumor-bearing C57BL/6J mice at 24, 72, and 168 hours pi. Mean $^{125}$I/$^{111}$In-ratio, standard deviation, and sample size per tissue are shown (mean ± sd (n=x)).

| **24 hours pi** | **CD3xTRP1** | **CD3xMock** | **TRP1xMock** |
|---|---|---|---|
| | $^{125}$I/$^{111}$In-ratio | $^{125}$I/$^{111}$In-ratio | $^{125}$I/$^{111}$In-ratio |
| **Blood** | 0.82 ± 0.02 (n=5) | 0.79 ± 0.01 (n=7) | 0.67 ± 0.02 (n=6) |
| **Skin (pigmented)** | 0.80 ± 0.00 (n=1) | 0.74 ± 0.04 (n=5) | 0.66 ± 0.02 (n=3) |
| **Skin (non-pigmented)** | 0.80 ± 0.01 (n=5) | 0.74 ± 0.02 (n=7) | 0.63 ± 0.04 (n=6) |
| **Brown adipose tissue** | 0.84 ± 0.02 (n=5) | 0.79 ± 0.03 (n=7) | 0.66 ± 0.02 (n=6) |
| **Lymph nodes** | 0.27 ± 0.04 (n=5) | 0.25 ± 0.03 (n=7) | 0.41 ± 0.05 (n=6) |
| **KPC3** | 0.71 ± 0.01 (n=4) | 0.66 ± 0.02 (n=7) | 0.56 ± 0.01 (n=5) |
| **KPC3-TRP1** | 0.45 ± 0.02 (n=5) | 0.66 ± 0.01 (n=6) | 0.43 ± 0.02 (n=6) |
| **Muscle** | 0.82 ± 0.00 (n=5) | 0.80 ± 0.02 (n=7) | 0.64 ± 0.03 (n=6) |
| **Thymus** | 0.45 ± 0.05 (n=5) | 0.44 ± 0.05 (n=7) | 0.64 ± 0.01 (n=6) |
| **Heart** | 0.82 ± 0.02 (n=5) | 0.79 ± 0.01 (n=7) | 0.66 ± 0.02 (n=6) |
| **Lung** | 0.80 ± 0.02 (n=5) | 0.75 ± 0.02 (n=7) | 0.65 ± 0.02 (n=6) |
| **Spleen** | 0.20 ± 0.01 (n=5) | 0.16 ± 0.01 (n=7) | 0.52 ± 0.02 (n=6) |
| **Pancreas** | 0.66 ± 0.02 (n=5) | 0.65 ± 0.03 (n=7) | 0.59 ± 0.02 (n=6) |
| **Stomach** | 0.84 ± 0.30 (n=5) | 0.77 ± 0.27 (n=7) | 0.73 ± 0.04 (n=6) |
| **Duodenum** | 0.98 ± 0.75 (n=5) | 0.75 ± 0.37 (n=7) | 0.64 ± 0.02 (n=6) |
| **Colon** | 0.71 ± 0.04 (n=5) | 0.68 ± 0.03 (n=7) | 0.61 ± 0.03 (n=6) |
| **Liver** | 0.63 ± 0.02 (n=5) | 0.60 ± 0.04 (n=7) | 0.59 ± 0.02 (n=6) |
| **Kidney** | 0.60 ± 0.02 (n=5) | 0.61 ± 0.02 (n=7) | 0.50 ± 0.01 (n=6) |
| **Prostate** | 0.85 ± 0.06 (n=5) | 0.78 ± 0.03 (n=7) | 0.65 ± 0.02 (n=6) |
| **Bone marrow** | 0.59 ± 0.04 (n=5) | 0.50 ± 0.03 (n=7) | 0.57 ± 0.04 (n=6) |
| **Bone** | 0.64 ± 0.02 (n=5) | 0.58 ± 0.02 (n=7) | 0.59 ± 0.03 (n=6) |

| 72 hours pi | CD3xTRP1 | CD3xMock | TRP1xMock |
|---|---|---|---|
| | $^{125}I/^{111}In$-ratio | $^{125}I/^{111}In$-ratio | $^{125}I/^{111}In$-ratio |
| **Blood** | 0.84 ± 0.02 (n=5) | 0.81 ± 0.03 (n=7) | 0.67 ± 0.01 (n=6) |
| **Skin (pigmented)** | 0.68 ± 0.01 (n=2) | 0.74 ± 0.06 (n=5) | 0.59 ± 0.02 (n=2) |
| **Skin (non-pigmented)** | 0.70 ± 0.10 (n=5) | 0.76 ± 0.06 (n=7) | 0.60 ± 0.03 (n=6) |
| **Brown adipose tissue** | 0.81 ± 0.03 (n=5) | 0.76 ± 0.07 (n=7) | 0.64 ± 0.01 (n=6) |
| **Lymph nodes** | 0.15 ± 0.00 (n=5) | 0.13 ± 0.02 (n=7) | 0.29 ± 0.05 (n=6) |
| **KPC3** | 0.62 ± 0.04 (n=5) | 0.60 ± 0.08 (n=7) | 0.50 ± 0.03 (n=6) |
| **KPC3-TRP1** | 0.22 ± 0.03 (n=5) | 0.53 ± 0.11 (n=7) | 0.27 ± 0.04 (n=6) |
| **Muscle** | 0.85 ± 0.05 (n=5) | 0.80 ± 0.07 (n=7) | 0.65 ± 0.02 (n=6) |
| **Thymus** | 0.25 ± 0.04 (n=5) | 0.24 ± 0.06 (n=7) | 0.53 ± 0.04 (n=6) |
| **Heart** | 0.81 ± 0.01 (n=5) | 0.80 ± 0.06 (n=7) | 0.65 ± 0.02 (n=6) |
| **Lung** | 0.76 ± 0.03 (n=5) | 0.74 ± 0.08 (n=7) | 0.63 ± 0.01 (n=6) |
| **Spleen** | 0.14 ± 0.02 (n=5) | 0.10 ± 0.03 (n=7) | 0.37 ± 0.02 (n=6) |
| **Pancreas** | 0.68 ± 0.17 (n=5) | 0.60 ± 0.03 (n=7) | 0.51 ± 0.03 (n=6) |
| **Stomach** | 0.85 ± 0.14 (n=5) | 0.78 ± 0.08 (n=7) | 0.72 ± 0.03 (n=6) |
| **Duodenum** | 0.48 ± 0.04 (n=5) | 0.43 ± 0.06 (n=7) | 0.57 ± 0.04 (n=6) |
| **Colon** | 0.69 ± 0.03 (n=5) | 0.61 ± 0.06 (n=7) | 0.59 ± 0.02 (n=6) |
| **Liver** | 0.52 ± 0.04 (n=5) | 0.59 ± 0.33 (n=7) | 0.51 ± 0.02 (n=6) |
| **Kidney** | 0.61 ± 0.05 (n=5) | 0.54 ± 0.09 (n=7) | 0.48 ± 0.02 (n=6) |
| **Prostate** | 0.78 ± 0.13 (n=5) | 0.76 ± 0.07 (n=7) | 0.60 ± 0.04 (n=6) |
| **Bone marrow** | 0.48 ± 0.04 (n=5) | 0.45 ± 0.08 (n=7) | 0.50 ± 0.05 (n=6) |
| **Bone** | 0.58 ± 0.03 (n=5) | 0.52 ± 0.05 (n=7) | 0.49 ± 0.03 (n=6) |

| 168 hours pi | CD3xTRP1 | CD3xMock | TRP1xMock |
|---|---|---|---|
| | $^{125}I/^{111}In$-ratio | $^{125}I/^{111}In$-ratio | $^{125}I/^{111}In$-ratio |
| **Blood** | 0.84 ± 0.03 (n=6) | 0.79 ± 0.03 (n=5) | 0.64 ± 0.01 (n=6) |
| **Skin (pigmented)** | 0.67 ± 0.05 (n=5) | 0.68 ± 0.04 (n=3) | 0.51 ± 0.04 (n=6) |
| **Skin (non-pigmented)** | 0.63 ± 0.09 (n=6) | 0.65 ± 0.07 (n=5) | 0.50 ± 0.04 (n=7) |
| **Brown adipose tissue** | 0.76 ± 0.12 (n=6) | 0.66 ± 0.06 (n=5) | 0.55 ± 0.03 (n=7) |
| **Lymph nodes** | 0.09 ± 0.02 (n=6) | 0.10 ± 0.02 (n=5) | 0.14 ± 0.02 (n=7) |
| **KPC3** | 0.57 ± 0.04 (n=6) | 0.51 ± 0.03 (n=4) | 0.39 ± 0.03 (n=7) |
| **KPC3-TRP1** | 0.21 ± 0.03 (n=6) | 0.55 ± 0.06 (n=5) | 0.20 ± 0.02 (n=7) |
| **Muscle** | 0.74 ± 0.04 (n=6) | 0.79 ± 0.09 (n=5) | 0.57 ± 0.01 (n=7) |
| **Thymus** | 0.19 ± 0.06 (n=6) | 0.16 ± 0.03 (n=5) | 0.37 ± 0.05 (n=7) |
| **Heart** | 0.80 ± 0.03 (n=6) | 0.74 ± 0.04 (n=5) | 0.60 ± 0.02 (n=7) |
| **Lung** | 0.73 ± 0.05 (n=6) | 0.65 ± 0.04 (n=5) | 0.54 ± 0.01 (n=7) |
| **Spleen** | 0.10 ± 0.01 (n=6) | 0.07 ± 0.00 (n=5) | 0.22 ± 0.02 (n=7) |
| **Pancreas** | 0.50 ± 0.07 (n=6) | 0.54 ± 0.09 (n=5) | 0.47 ± 0.02 (n=7) |
| **Stomach** | 0.90 ± 0.10 (n=6) | 1.10 ± 0.53 (n=5) | 0.81 ± 0.14 (n=7) |
| **Duodenum** | 0.38 ± 0.03 (n=6) | 0.31 ± 0.02 (n=5) | 0.48 ± 0.03 (n=7) |
| **Colon** | 0.59 ± 0.07 (n=6) | 0.54 ± 0.05 (n=5) | 0.50 ± 0.04 (n=7) |
| **Liver** | 0.39 ± 0.07 (n=6) | 0.33 ± 0.02 (n=5) | 0.37 ± 0.03 (n=7) |
| **Kidney** | 0.62 ± 0.18 (n=6) | 0.56 ± 0.02 (n=5) | 0.39 ± 0.03 (n=7) |
| **Prostate** | 0.77 ± 0.14 (n=6) | 0.65 ± 0.06 (n=5) | 0.55 ± 0.06 (n=7) |
| **Bone marrow** | 0.35 ± 0.07 (n=6) | 0.26 ± 0.06 (n=5) | 0.28 ± 0.04 (n=7) |
| **Bone** | 0.43 ± 0.03 (n=6) | 0.36 ± 0.03 (n=5) | 0.34 ± 0.02 (n=7) |

**Table S5. Serum fraction of blood circulating [$^{111}$In]In-CD3xTRP1 and control bsAbs over time.** Serum fraction of blood circulating [$^{111}$In]In-CD3xTRP1, [$^{111}$In]In-CD3xMock, and [$^{111}$In]In-TRP1xMock in C57BL/6J mice bearing KPC3-TRP1 and KPC3 tumors that received 12.5 µg [$^{111}$In]In-bsAb intraperitoneally at 24, 72, and 168 hours pi. Mean $^{111}$In serum fraction, standard deviation, and sample size are shown (mean ± sd (n=x)).

| | **24 hours pi** | **72 hours pi** | **168 hours pi** |
|---|---|---|---|
| | (%) | (%) | (%) |
| **CD3xTRP1** | 75.4 ± 5.4 (n=5) | 73.4 ± 3.7 (n=5) | 80.7 ± 5.9 (n=6) |
| **CD3xMock** | 76.8 ± 5.1 (n=6) | 80.6 ± 6.9 (n=7) | 81.0 ± 7.4 (n=5) |
| **TRP1xMock** | 79.5 ± 4.3 (n=6) | 77.6 ± 3.5 (n=6) | 78.3 ± 6.0 (n=7) |
